# Structural insights into the mechanism of the sodium/iodide symporter (NIS)

**DOI:** 10.1101/2022.04.07.487502

**Authors:** Silvia Ravera, Juan Pablo Nicola, Glicella Salazar de Simone, Fred J. Sigworth, Erkan Karakas, L. Mario Amzel, Mario A. Bianchet, Nancy Carrasco

## Abstract

The sodium/iodide symporter (NIS) is the essential plasma membrane protein that mediates active iodide (I^-^) transport into the thyroid gland, the first step in the biosynthesis of the thyroid hormones—the master regulators of intermediary metabolism. NIS couples the inward translocation of I^-^ against its electrochemical gradient to the inward transport of Na^+^ down its electrochemical gradient. For nearly 50 years before its molecular identification, NIS was already the molecule at the center of the single most effective internal radiation cancer therapy ever devised: radioiodide (^131^I^-^) treatment for thyroid cancer. Mutations in NIS cause congenital hypothyroidism, which must be treated immediately after birth to prevent stunted growth and cognitive deficiency. To date, the structure of NIS has been unknown. Here, we report three structures of rat NIS, determined by single-particle cryo-electron microscopy (cryo-EM): one with no substrates bound, one with 2 Na^+^ and 1 I^-^ bound, and one with 1 Na^+^ and the oxyanion perrhenate bound. Structural analyses, functional characterization, and computational studies reveal the substrate binding sites and residues key for transport activity. Our results yield insights into how NIS selects, couples, and translocates anions—thereby establishing a framework for understanding NIS function—and into how it transports different substrates with different stoichiometries and releases substrates from its substrate-binding cavity into the cytosol.

---

Iodide (I^-^) is a key micronutrient because its oxidized form, iodine, is an essential constituent of the thyroid hormones (THs). An adequate supply of I^-^ early in life is crucial for preventing I^-^ deficiency disorders (IDDs), because the THs are critical for embryonic and postembryonic development, particularly that of the central nervous system, lungs, and musculoskeletal system. The THs are also master regulators of cellular metabolism in virtually all tissues at all stages of life^1,2^. The first step in the biosynthesis of the THs is the active transport of I^-^ into the thyroid, which is mediated by the product of the SLC5A5 gene: the sodium/iodide symporter (NIS), an intrinsic plasma membrane protein located at the basolateral surface of the thyroid follicular cells. NIS couples the inward translocation of I^-^ against its electrochemical gradient to the inward transport of Na^+^ down its electrochemical gradient^3^. One of the most remarkable properties of NIS is that—unlike Cl^-^ channels and transporters, which transport both Cl^-^ and I^-^ (with affinities in the mM range)^4^—NIS discriminates between these two anions in an astonishingly exquisite way that is of great physiological significance: it transports I^-^ but not Cl^-^, even though the concentration of Cl^-^ in the extracellular milieu (∼100 mM) is over 10^5^ times that of I^-^ (<1 μM). Significant mechanistic insights have been gained by determining how NIS mutations found in patients with IDDs affect the protein’s folding, plasma membrane targeting, and activity^5,6^.

NIS is also the molecule at the center of the remarkably successful treatment for thyroid cancer based on ^131^I^-^, administered after thyroidectomy: this therapy targets remnant malignant cells and metastases that actively accumulate the radioisotope via NIS. In addition, NIS is functionally expressed endogenously not only in several extrathyroidal tissues—such as the salivary glands, stomach, intestine, and lactating breast—but also in primary and metastatic breast cancers^7^. This last finding opens up the possibility that NIS-mediated ^131^I^-^ treatment could be used against breast cancer. Furthermore, there is great interest^5^ in expressing NIS exogenously in cancers that do not express it endogenously to render them susceptible to destruction by ^131^I^-^ (^refs. 8,9^).

NIS translocates several substrates other than I^-^, including oxyanions such as the environmental pollutant perchlorate (ClO_4_^-^), and the imaging substrates pertechnetate (^99m^TcO_4_^-^) and perrhenate (^186^ReO_4_^-, 188^ReO_4_^-^), which are used in single-photon emission computed tomography (SPECT). The NIS substrates tetrafluoroborate (^18^F-BF_4_^-^) and ^124^I^-^ are used in positron emission tomography (PET). NIS is becoming the counterpart of green fluorescent protein (GFP) or luciferase for imaging studies in humans^5,8,10,11^. Although the physiology, biochemistry, biophysics, and cell biology of NIS have all been investigated extensively^1,5^, our understanding of its symport mechanism has, until now, been hindered by a lack of high-resolution structures of NIS. Here, we use single-particle cryo-electron microscopy (cryo-EM), molecular dynamics (MD) simulations, and functional studies to gain a structural understanding of how NIS binds its anion substrates with such high affinity, how it translocates anions, and how mutations in NIS lead to congenital IDDs.

## Results

### Structure determination

Rat NIS shares 89% sequence identity with human NIS (Supplementary Fig. 1) and has similar transport properties^6,12-14^. We have previously shown that glycosylation is not required for NIS activity^12,15^. Hence, to carry out structural studies, we engineered a cDNA coding for a recombinant unglycosylated full-length rat NIS molecule: a triple NIS mutant in which all three Asn residues that are glycosylated were replaced with Gln (N225Q/N485Q/N497Q), with an HA (human influenza hemagglutinin) tag at the N-terminus and HIS and SBP (streptavidin-binding protein) tags at the C-terminus (recombinant tagged NIS, rtNIS). We expressed rtNIS in 293F cells, enriched for cells expressing the protein by flow cytometry with an anti-HA antibody, and demonstrated that its activity is indistinguishable from that of WT NIS (Extended Data Fig. 1 a-c). We then solubilized and purified rtNIS. The protein that eluted in the peak fraction and electrophoresed as a single polypeptide was used to determine the structure of NIS by single-particle cryo-EM (Extended Data Table 1, Extended Data Fig. 1 d-k).

The cryo-EM map (overall resolution 3.46 Å) reveals that NIS in the micelles assumes a dimeric conformation with excellent density for all 26 transmembrane segments (TMSs) as well as for significant regions of the extramembrane loops and 17 out of 74 residues in the C-terminus. Well-resolved TMSs of different lengths displayed clearly visible α-helical features (Extended Data Fig. 2a). As a result, the map enabled us to build a model of the entire protein from residue 9 to residue 561, with the exception of the loops between TMSs 1 and 2 (residues 34-52), TMSs 5 and 6 (residues 183-188), and TMSs 12 and 13 (residues 483-510), whose lack of density may be attributable to the high flexibility of these regions. To further improve the quality of the map, we applied a symmetry expansion to the particles and carried out a local refinement with a mask designed for a single monomer yielding a new map at 3.30 Å resolution (Extended Data Fig. 1k).

The NIS dimers embedded in each micelle are in an antiparallel configuration (Fig. 1a, b). As we have demonstrated, when NIS is endogenously expressed, its N-terminus faces the extracellular milieu and its C-terminus the cytosol^12,15-18^. Thus, it is extremely unlikely that the antiparallel NIS dimers in the micelles are physiological. Instead, the antiparallel configuration resulted from the solubilization and purification process. The two oppositely oriented monomers interact through a two-fold axis perpendicular to TMS 5 and TMS 10 and to their local symmetry counterparts. No salt bridges are observed at the interface, and there are only two symmetrical pairs of polar residues across the local symmetry axis—S396 and T400—but they do not appear to contribute to forming the dimer by means of hydrogen bonds or water-mediated interactions. Only apolar interactions hold the dimer together. Duplicated by the local symmetry, aromatic stacking of W165 between F385 and F392 in the other monomer allows for a specific interaction required to generate identical dimers in different micelles (Extended Data Fig. 2b).

**Fig. 1.**
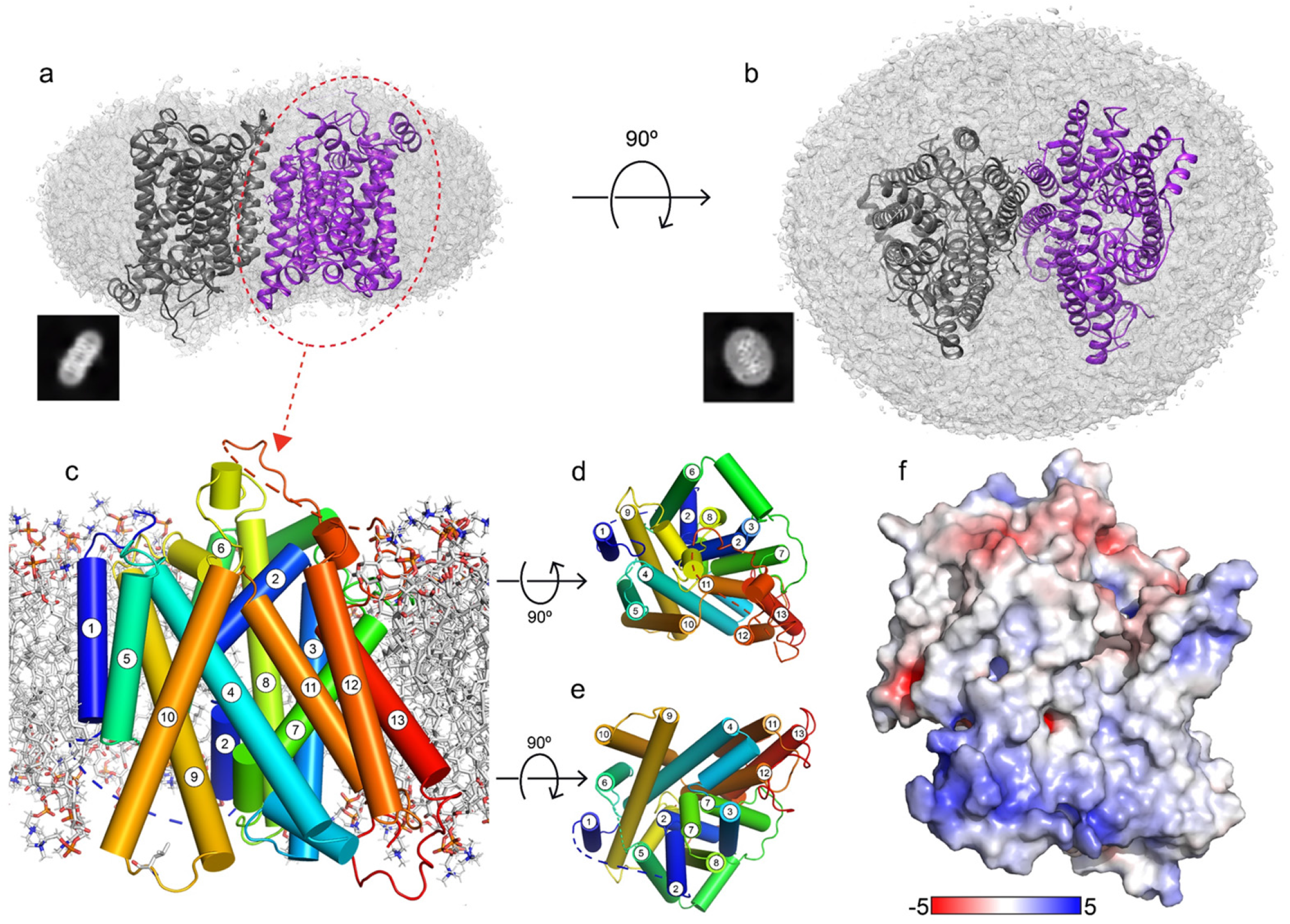
Structure of NIS. **a-b**. Two NIS molecules entirely embedded in a detergent micelle. **a**. Side view. **b**. Top view. Examples of 2D classes representing the corresponding views are shown in the black squares.) **c**. NIS structure viewed from the plane of the membrane, with the numbered TM helices depicted as cylinders; **d**, the corresponding top view, and **e**, bottom view. **f**. Side view of NIS as in (c) but with solvent-accessible surface colored according to the electrostatic potential in k_B_T/e (red, negative; blue, positive).

As predicted^14,19^, NIS has a LeuT fold with two 5-helix bundle domains (TMSs 2-6 and 7-11) related by an internal pseudo-two-fold symmetry around an axis through the center of the transporter and perpendicular to the plane of the membrane (Fig. 1c, Extended Data Fig. 1a, Supplementary Fig. 2). This conserved molecular architecture is common to various membrane transporters that have unrelated sequences and belong to different families^20-24^. The alignment between the two repeats has a root mean square deviation (RMSD) of 3.01 Å for 115 Cα atoms aligned [LeuT RMSD 5.3 Å for 130 Cα atoms^25^]. TMSs 2-6 are related to TMSs 7-11 by a 178° rotation parallel to the membrane plane. TMSs 2 and 7 have unwound regions (S66-A70 and G261-A263), which likely bind ions and/or participate in the conformational changes that occur during the transport cycle. An analysis of the secondary structure motifs revealed the presence of π-helices at three positions: 101-105, 247-251, and 366-371. Although the protein was purified in the presence of a high concentration of Na^+^ (350 mM), we refer to this structure as apo-NIS, as it does not contain any densities attributable to bound ions.

The electrostatic potential surface shows that the distribution of charges satisfies the positive-inside rule of transmembrane protein topology (Fig. 1f). NIS adopts an inward-facing conformation similar to those observed for the *Vibrio cholerae* Na^+^/galactose transporter (vSGLT)^26^ and human Na^+^/glucose transporter 1 (hSGLT1)^24^, and opposite that of the outwardly open hSGLT2^23^ (Supplementary Fig. 3) (Cα RMSDs = 2.9 Å over 313 residues, 3.3 Å over 378 residues, and 3.75 Å over 363 residues, respectively). Although the positions of the TMSs of NIS suggest that the conformation is inwardly open, the opening is narrower than those in vSGLT or hSGLT1.

### Iodide and sodium binding sites

To elucidate how NIS binds I^-^ with such high affinity, we determined the structure of NIS with I^-^ bound to it at 3.12 Å (Extended Data Fig. 3a). Even though the NIS-I^-^ structure is highly similar to that of apo-NIS (RMSD = 0.86 over 495 residues), we observed three additional non-protein densities in the core of the NIS-I^-^ map that correspond to the substrates. To assign the correct ion to each position, we plotted the density map in the vicinity of each ion site and generated a sphere with a radius of 5 Å from the density center. Starting at an ion position, the map density (which is proportional to the electrostatic potential) was ascertained along 48 radial lines in uniformly distributed directions. The radial density, given as the mean over all angles, is plotted as a black curve (Fig. 2a). The height and width of the peaks are expected to reflect the size of the ions: the positive central peak tends to reflect the nuclear charge, and the width of the peak corresponds to the spatial extent of the ion’s electron density^27^. The undershoot reflects the potential outside the valence electrons and will be more negative for an anion. Thus, we conclude that the largest peak (1.5 arbitrary units, AU) corresponds to I^-^, and the smaller peaks (1.0 and 1.2 AU) to Na^+^ (Fig. 2a).

**Fig. 2.**
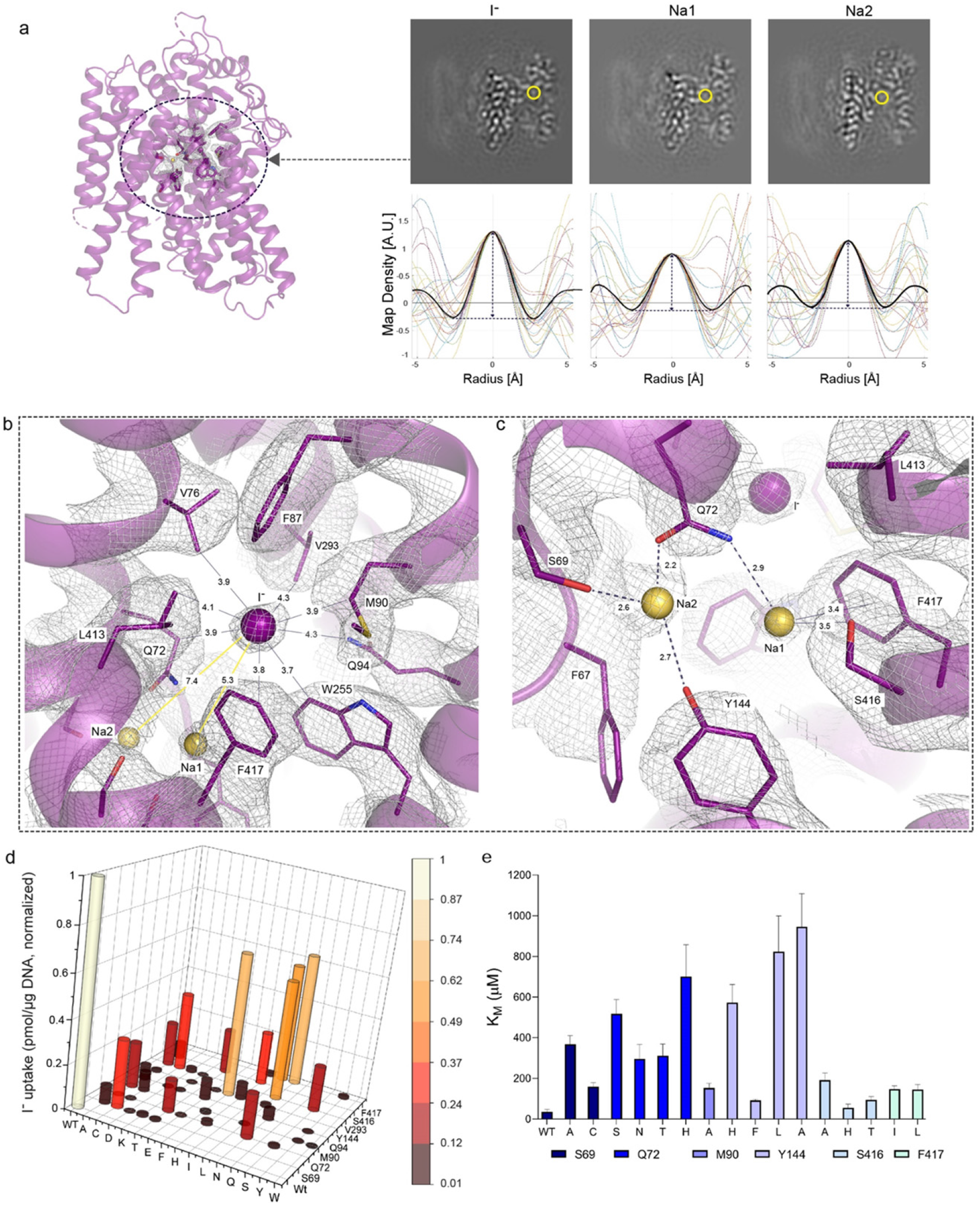
Iodide and sodium binding sites. Fig. 2 | Iodide and sodium binding sites. **a**. In the NIS-I^-^ structure, the yellow circle indicates the substrate binding pocket. Ions transported by NIS were identified by analyzing the map density values (map slices in upper panels) along 24 lines passing through each site, circled in yellow. Map values are plotted for each line (lower panels), and the spherically-averaged mean is plotted in black. **b**. Close-up of the I^-^ binding pocket, where I^-^ is represented by the purple sphere. The distance (in Å) from I^-^ to each surrounding residue is indicated. Yellow spheres represent sodium ions. **c**. Close-up of interactions between Na1 and Na2 and surrounding residues. **d**. Functional effects of mutations in binding-site residues. Steady-state transport was determined at 20 μM I^-^ and 140 mM Na^+^ for 30 min, normalized to values obtained with WT NIS (values obtained in the presence of ClO_4_^-^, which are < 10% of the values obtained in its absence, have already been subtracted). **e**. K_M_ values obtained from the kinetic analysis of I^-^ transport.

The I^-^ binding site is delimited by residues Q72, V76, M90, Q94, W255, V293, L413, and F417 (Fig. 2b), and has a positive residual charge that further stabilizes the anion (Extended Data Fig. 4). All these residues are 100% conserved from humans to fish (Supplementary Fig. 1); notably, most of them are hydrophobic.

In addition to the density in NIS-I^-^ ascribed to I^-^, two other clear densities were present, which we attribut to Na^+^ ions. One of them, which we dubbed Na1 because it is the closer one to the I^-^ (5.3 Å), interacts with Q72 and S416, and is stabilized by a cation-π interaction with the aromatic ring of F417 (Fig. 2c). The other Na^+^, by contrast—which we named Na2 (7.4 Å from I^-^) —is coordinated by the side chain oxygen atoms of three residues: S69, Q72, and Y144 (Fig. 2c).

To ascertain the functional relevance of the interactions observed, we engineered several amino acid substitutions at positions 69, 72, 90, 94, 144, 293, 416, and 417. In all cases, I^-^ transport at steady state (at 20 μM I^-^) was substantially reduced, and no I^-^ was transported at all when the substituted residues were charged (Fig. 2d). Similar results have been reported with substitutions at some of these positions in human NIS^28^. Here we show that the effects of the replacements were clearly not due to impaired NIS mutant protein expression or trafficking to the cell surface, because all mutants were present at the plasma membrane at levels comparable to those of WT NIS, as determined by flow cytometry with an antibody against the extracellularly facing HA tag (Extended Data Fig. 1a, Supplementary Fig. 4). Increasing the concentration of I^-^ 10-fold (to 200 μM I^-^) caused I^-^ accumulation by WT NIS to increase 2- to 3-fold, but had a more pronounced effect on the mutants, whose I^-^ accumulation increased 3- to 20-fold (Extended Data Fig. 5), suggesting that the mutants had a markedly lower affinity for I^-^ than WT NIS. To test this hypothesis, we carried out I^-^ transport assays at initial rates. Indeed, the K_M_ values of the mutants for I^-^ were considerably higher (3–30-fold) than that of WT NIS (20-30 μM) (Fig. 2e, Extended Data Fig. 5). All mutant NIS proteins also had a higher K_M_ for Na^+^ than WT NIS (50 mM) (Extended Data Fig. 5). These data are consistent with the fact that the binding of one of the substrates increases the affinity of NIS for the other^29,30^.

### Oxyanion binding site

Although NIS discriminates finely between I^-^ and the other halides, it does translocate substrates other than I^-^, including the oxyanions ^99m^TcO_4_^-^, ReO_4_^-^, and ClO_4_^-^ (an environmental pollutant)^5,7,31-33^. Strikingly, whereas NIS transports I^-^ electrogenically (with a 2 Na^+^ : 1 I^-^ stoichiometry), it transports oxyanions electroneutrally (with a 1 Na^+^ : 1 oxyanion stoichiometry)^14,29,34^. To deepen our understanding of oxyanion transport by NIS, we determined the structure of the protein in complex with ReO_4_^-^ at a resolution of 3.15 Å (Extended Data Fig. 3b). We observed two densities located at approximately the same position as two of the three densities in the NIS-I^-^ map. We propose, based on an analysis similar to the one carried out for the NIS-I^-^ structure (Fig. 2a), that the large density corresponds to ReO_4_^-^, owing to the size (2 AU) and shape of the peak, and the small one (1 AU) to Na^+^ (Extended Data Fig. 6).

ReO_4_^-^ forms hydrogen bonds with Q72 (2.5 Å) and Q94 (2.8 Å), and it is in close contact (<3.6 Å) with V76, M90, W255, V293, and F417 (Fig. 3a). A Na^+^ is located in a very similar position to the one that Na1 occupies in the NIS-I^-^ structure, and it interacts with the same residues: Q72, S416, and F417 (Fig. 3a). No other density was found that could be attributed to a second Na^+^, which is consistent with the 1:1 stoichiometry of NIS-mediated oxyanion transport^7,14,29,31,33-35^. Although the charge center distance between the cation and the oxyanion is the same as that between I^-^ and Na1 (5.3 Å), the distance between Na1 and a ReO_4_^-^ oxygen atom is shorter (3.9 Å). That the amino acids shown to coordinate ReO_4_^-^ play a significant role in oxyanion transport was demonstrated by the effects of the amino acid substitutions on ReO _4_ transport (Extended Data Figs. 6 b, 7 a-d). A detailed analysis of the Q72 mutants showed that NIS with Ala, Cys, Glu, or His at this position does not transport ReO_4_^-^ at a concentration of 3 μM. However, these mutants do transport a modest amount of ReO_4_^-^ when its concentration is increased 10-fold. In agreement with these data, the K_M_ of Q72H NIS for ReO_4_^-^ was 10 times that of WT NIS (Extended Data Fig. 7 e,f).

**Fig. 3.**
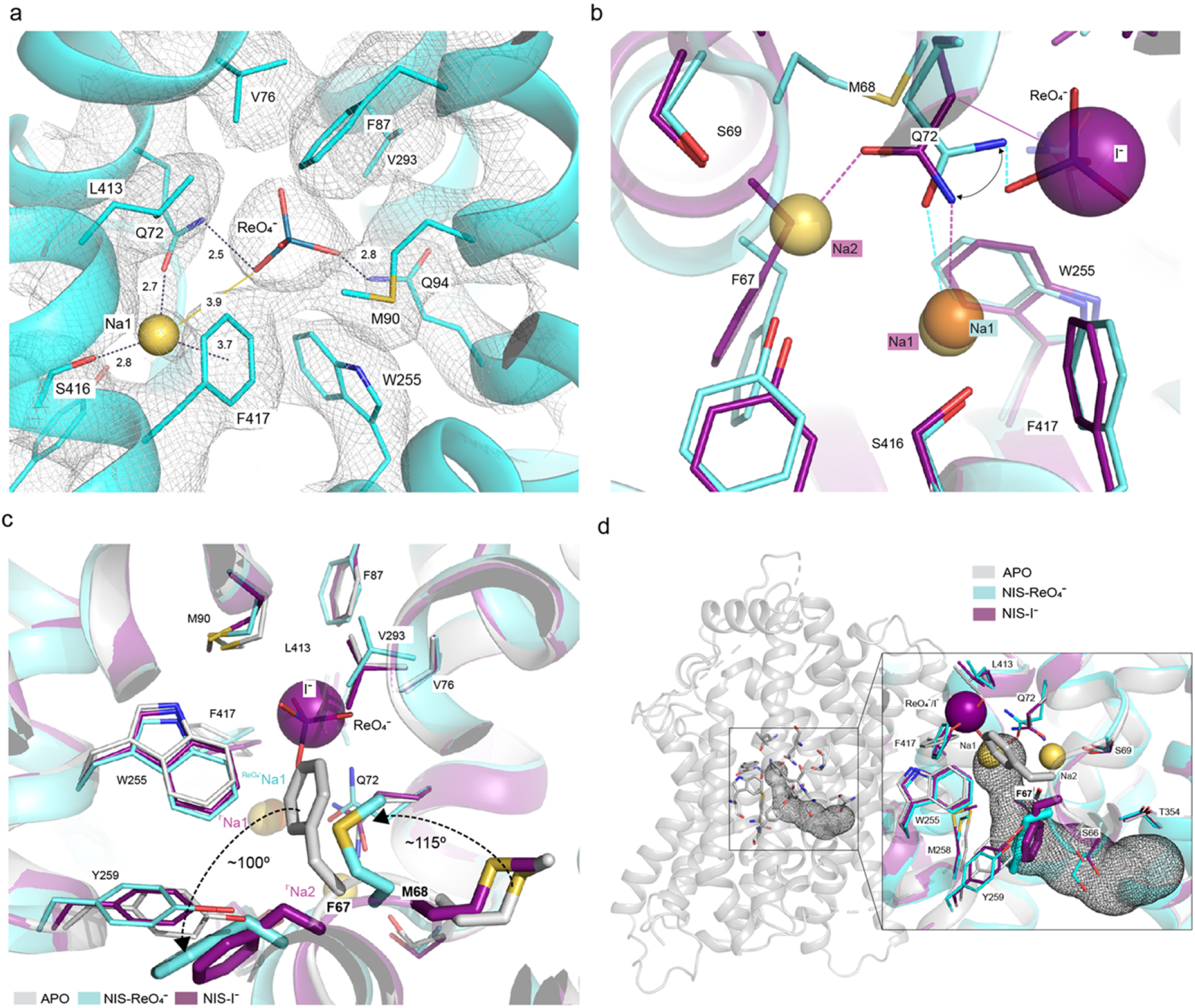
Perrhenate binding pocket and localized changes caused by ion binding. **a**. Close-up of the ReO_4_^-^ binding pocket. ReO_4_^-^ is represented by sticks and Na1 by a yellow sphere. Distances from ReO_4_^-^ and Na1 to surrounding residues (within 3.9 Å) are indicated, as is the distance from ReO_4_^-^ to Na1. **b**. Close-up of the structural alignment of NIS-I^-^ and NIS-ReO_4_^-^ showing the shift of Q72. **c**. Close-up of the structural alignment between apo-NIS, NIS-I^-^ and NIS-ReO_4_^-^. The arrows represent the shift of F67 and M68 from the position they occupy in the apo-NIS structure to the one in the substrate-bound structures. In the apo-NIS structure, F67 partially occupies the anion site. **d**. Tunnel calculated by Caver-Analyst in the apo-NIS structure that connects the substrate binding pocket with the cytosol. Inset: alignment of the three structures revealing that in the substrate-bound NIS, F67 blocks the pathway from the binding sites to the cytosol.

### Electrogenic versus electroneutral transport

The NIS-ReO_4_^-^ structure has a Gln rotamer at position 72 has a different rotamer (Fig. 3b) from the one in the NIS-I^-^ structure. As a result, Q72 interacts with ReO_4_^-^ through its amino group and with Na^+^ at the Na1 site through its carbonyl group (which in NIS-I^-^ interacts with Na^+^ at the Na2 site). This movement of Q72 might explain, at least in part, the absence of Na^+^ at the Na2 site in the NIS-ReO_4_^-^ structure and, as a consequence, the electroneutral stoichiometry of Na^+^/ReO_4_^-^ transport.

### Ion binding causes localized structural changes in NIS

The three NIS structures are remarkably similar. The small RMSDs between the apo and ion-bound structures (0.86 Å for NIS-I^-^ and 1.0 Å for NIS-ReO_4_^-^) for 495 Cα atoms aligned suggest that binding of the ions to NIS causes only small, localized changes. Of these structural changes, the most significant occur in the TMS2 kink between F66 and Q72. In the apo structure, this kink is stabilized by hydrogen bonds between the amino group of Q72 and the hydroxyl group of S69 and between the former and the main chain carbonyl of F67, as well as by a stacking interaction between the aliphatic portion of the side chain of Q72 and the aromatic ring of F67. The kink is further stabilized by a hydrogen bond formed between Y144, in the adjacent TMS4, and the main chain carbonyl of S66 (Fig. 4a).

**Fig. 4.**
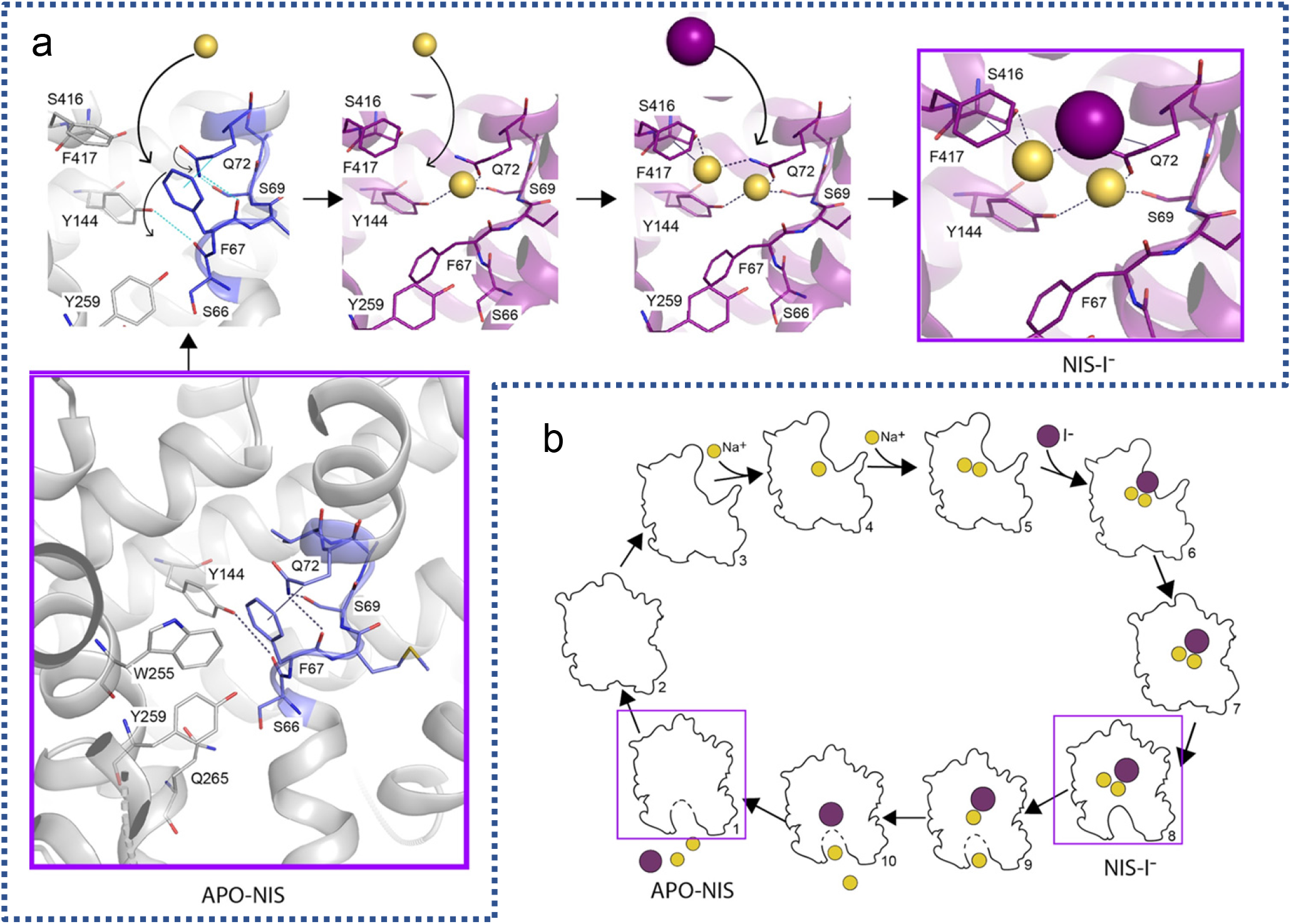
NIS mechanism. **a**. Mechanism of NIS substrate binding, starting from the apo-NIS structure. A hydrogen-bond network between F67, S69, Q72, and Y144 and a hydrophobic stacking interaction between F67 and Q72 are disrupted by the binding of the first Na^+^. This facilitates the binding of a second Na^+^ and subsequently of I^-^. **b**. Cartoon of the putative NIS transport cycle. The experimental structures are enclosed in magenta boxes.

In the NIS-I^-^ structure, these kink-stabilizing residues are rearranged to coordinate the Na^+^ ions. Q72 flips from its position in the apo conformation to coordinate Na2 with its amide oxygen and Na1 with its amide nitrogen (Figs. 2b, c, 4a). The aromatic ring of F417 stabilizes Na1 through a cation-π interaction (Figs. 2c, 4a). It is possible that Na1, which appears to be coordinated to a lesser extent than Na2, is also stabilized by the negative charge of the anion. These rearrangements of the TMS2 kink make F67 swing 110° away from the inside of the anion binding pocket to the cytosolic side of the substrate site. In the NIS-ReO_4_^-^ structure the changes in the TMS2-kink are more pronounced. The backbone atoms of F67 also move 2 Å away from the center, and M68 rotates ∼115° upward from the position it has in both the apo-NIS and NIS-I^-^ structures (Fig. 3c). Given the position of the carbonyl of F67, and because amino acid substitutions at position 67 reduce I^-^ transport more than ReO_4_^-^ transport (Supplementary Fig. 5), we can speculate that the local rearrangements brought about by the replacements interfere with the coordination of Na2 and, consequently, ReO_4_^-^ transport should be less affected. Notably, other residues in the structures with substrates bound do not occupy significantly different positions than they do in the apo-NIS structure.

Using the Caver-Analyst software^36^, we identified in the apo-NIS structure a hydrophilic tunnel that connects the substrate-binding cavity to the cytosol (Fig. 3d). This tunnel is delimited by TMSs 2, 4, 7, and 9, and lined by some of the residues known to be important for NIS function^37^ and others reported to be mutated in patients with congenital hypothyroidism due to mutations in NIS. In the apo structure, the ring of F67 partially occupies the anion binding pocket. On the other hand, in the substrate-bound NIS structures, the substrate-binding cavity is cut off from the exit pathway by the side chain of F67, suggesting that the latter structures are in an occluded conformation (Fig. 3d). Similar tunnels exist in the substrate-bound structures when the side chain of F67 is not taken into account during the calculations.

We ran MD simulations with the NIS-I^-^ structure, restraining the cations bound to the site observed in the experimental structure. A pathway analysis carried out on ∼1000 snapshots sampled every 50 ns revealed a tunnel similar to the one in the apo-NIS structure in 86% of the snapshots. This suggests that this tunnel corresponds to a substrate-release pathway, and that the NIS-I^-^ structure is in an occluded conformation (Fig. 3d), but poised to open up into the cytosol (Supplementary Figs. 3 and 6). The cation restraint was sufficient for the I^-^ to remain bound for an aggregated sampling time of 5 μs, suggesting that the cations are released before I^-^ is.

F87 and L413 are at the ceiling of the binding cavity (Fig. 2b), and Q414 (which is flexible and hydrophilic) is just above the cavity. These residues visited a number of conformers during the MD simulations (Extended Data Fig. 8). The side chains of the conformers swing up to 100°, thus allowing the ions to enter the symporter from the extracellular milieu. In MD simulations with NIS-I^-^, F67 fluctuates between conformations that open up the ion pocket to the exit path and conformations that interact with the anion (Supplementary Fig. 7). The latter interactions may help the ions exit.

## Discussion

In view of all these results, we propose that the structures we obtained correspond to the ones boxed in Fig. 4 a,b. We have shown—on the basis of state occupancies derived from kinetic data analyzed by statistical thermodynamics^29^—that, at physiological concentrations of Na^+^, 79% of NIS molecules have two Na^+^ ions bound to them, which increases the affinity of NIS for I^-^ by a factor of 10 (Fig. 4b, panels 5-6). We hypothesize that the ternary complex isomerizes, causing NIS to adopt an occluded conformation that poises it to open up into the cytoplasm and release its substrates (Fig. 4b, panels 7 to 1), and the unloaded symporter isomerizes to bind ions from the extracellular milieu and begin the next transport cycle (Fig. 4b, panels 2-4). The transport cycle of NIS, then, includes minimally the steps just described.

An essential part of the mechanism of NIS is the coupling of Na^+^ transport to I^-^ transport. The NIS structures reported here reveal that, when Na^+^ binds to the Na2 site, it disrupts the hydrogen-bond network that stabilizes the TMS2 kink in the apo-NIS structure (Fig. 4a), causing a conformational change that pushes F67 away from the anion binding site (Fig. 3c, d), thereby increasing the affinity of NIS for Na^+^ at the Na1 site and for I^-^. In the NIS-ReO_4_^-^ structure, the oxyanion interacts with Q72, moving it away from the Na2 site and thereby preventing it from coordinating Na^+^ at this site. In both I^-^ and oxyanion transport, the binding of Na^+^ at the Na2 site can trigger the structural change that enables the anion to bind (Extended Data Fig. 9). An alternative mechanism in which anion binding drives the conformational changes seems unlikely, given that NIS has a low affinity for the anion in the absence of Na^+^ (^ref.29^).

How does NIS manage to stabilize so many ions in such close proximity to one another in the NIS-I^-^ structure? A possible answer is that the cation reduces the pKa of Q72, deprotonating the amino group and generating a negative charge at the carbonyl of the amide group, thus compensating for the positive charge of Na^+^ at the Na2 site and helping to bind Na^+^ at the Na1 site, with the electron pair remaining in the amide. Because I^-^ is an extremely weak base, the base required for the proposed electron rearrangement might be provided by solvent, i.e., water molecules (or OH^-^) that may access the cation sites from the cytosolic side.

One of the most striking attributes of NIS is its ability to transport I^-^, a very scarce ion, at the submicromolar concentrations at which it is found in the extracellular milieu, as it is difficult to bind a halide ion such as I^-^ with high affinity. There remains, however, a very intriguing open question: Why is the affinity of NIS for I^-^ so high? The answer lies in the special properties of I^-^. As a 4d^10^5s^2^5p^6^ moiety (identical to xenon), I^-^ is surrounded by a large unreactive electron cloud that endows it with unique properties that have been elucidated in previous computational studies. Sun *et al*. ^38^ investigated the behavior of Na^+^ halides in a system with a water/air interface and found that I^-^ behaves differently than do the other halides: not only does it show the highest number-density close to the interface, but also its radial distribution (ρ(r)) has its maximum several Å closer to the water/air interface than those of the other halides. These observations strongly suggest that, given the opportunity, I^-^ will interact with a hydrophobic environment. In agreement with this conclusion is our experimental evidence that most of the residues surrounding I^-^ are hydrophobic, and replacing them with polar residues affects NIS activity (Fig. 2b, d-e, Extended Fig. 5). Furhermore, the ability of NIS to discriminate so exquisitely between Cl^-^ and I^-^ seems to be due to both the size and the hydrophobicity of I^-^ (^refs. 39-41^).

That nature conserved the same structural fold across proteins that transport chemically diverse substrates may indicate that thousands of years of evolution optimized a way to harness the energy stored in the Na^+^ electrochemical gradient to transport hydrophilic substrates against their concentration or electrochemical gradients—across the hydrophobic environment of the plasma membrane. To date, the prevailing expectation about a wide variety of mammalian transporters (SERT^22^, NaCT^21^, NKCC1^20^, SGLT1^24^, SGLT2^23^, SMCT^24^ etc.) has been that the location of their Na2 binding site will correspond to that of the well-known canonical Na2 binding site in the LeuT structure. This was indeed the case in SERT^22,42,43^, a member of the SLC6 family, although it is the only structure of a mammalian transporter in any of whose structures Na^+^ has been observed.

The structure of NIS with Na^+^ ions bound that we are reporting here is the first structure of a member of the SLC5 family in which bound Na^+^ ions have been visualized. The Na^+^ ions bound to NIS are in the ion binding cavity. Remarkably, the canonical hydroxyl group–containing residues (S353 and T354), though important for NIS function^19,37^, are in the exit pathway, ***not*** in the binding pocket. This is not entirely surprising, given how different I^-^ is from leucine or serotonin, but it does defy the expectation that the Na2 site should correspond to the canonical site in LeuT. Therefore, it will be of great interest to determine whether Na^+^ ions bind to other members of the SLC5 family and other families at the same site as they do in NIS, at other sites, or at the LeuT canonical site. The members of the SLC5 family, in particular, transport carbohydrates (glucose, galactose, myoinositol), organic cations (choline), and monocarboxylates (nicotinate, butyrate, etc.); among all these substrates, I^-^ is the only halide anion^23,24^. Although all these proteins have (or are predicted to have) the same fold, it will not be surprising if critical differences are found when their structures are determined, given that they transport extremely different substrates. Alignment of the sequences of the members of the SLC5 family (Supplementary Fig. 8) shows that certain residues in NIS are unique. For example, Q72 interacts with all three substrates in the NIS-I^-^ structure, but the equivalent residue is a His in the sugar transporters and an Ala in the choline transporter. Q72H NIS and Q72A NIS are inactive, highlighting the critical role that Q72 plays in NIS function (Fig. 2d).

NIS is of great significance in the basic field of plasma membrane transporters and for translational applications ranging from imaging studies to cancer treatment. In addition, NIS has been increasingly important as a reporter molecule in pre-clinical and clinical gene transfer studies, making it possible to monitor the delivery of therapeutic genes and oncolytic viruses^5,8,9^. There is consequently no question that the determination of the NIS structures reported here will have a profound impact. The structures have not only greatly deepened our understanding of the NIS mechanism and transport cycle, but also enabled us to elucidate the roles of some specific amino acids mutated in ITD patients in NIS function and targeting to the plasma membrane^5,6^ (Extended Data Fig. 10). The results presented here will pave the way for future determinations of the structure of NIS in different conformations and NIS molecules with different substrate selectivities and different stoichiometries, which will likely be extremely valuable in gene transfer studies and thus broaden the range of clinical applications of NIS.

## Data availability

Cryo-EM maps and atomic coordinates of the structures presented in this manuscript have been deposited in the PDB and Electron Microscopy Data Bank under the ID codes 7UUY and EMD-26806 for Apo-NIS; 7UV0 and EMD-26808 for NIS-I^-^; and 7UUZ and EMD-26807 for NIS-ReO_4_^-^, (pending approval).

## Acknowledgements

We thank Dr. Hassane Mchaourab and the members of the Carrasco laboratory for critical reading of the manuscript and insightful discussion. This study was supported by National Institutes of Health grants GM R01 114250 (to NC and LMA), NS021501 (to FJS), and NINDS R21NS108842 and a Pamela Mars Wright Innovator Award (MAB). We are grateful to Shenping Wu of the Yale CryoEM Resource facility for screening and data collection. This work was conducted in part using the CPU and GPU resources of the Advanced Computing Center for Research and Education (ACCRE) at Vanderbilt University and at the Maryland Advanced Research Computing Center (MARCC). We used the DORS storage system supported by the U.S. National Institute of Health (NIH) (S10RR031634 to Jarrod Smith).

## Author contributions

S.R., N.C. and L.M.A. conceived the project. S.R. expressed and purified the protein, and prepared cryo-grids. S.R. and E.K. processed the cryo-EM data. S.R., E.K., L.M.A., and M.A.B. built and refined the atomic models. F.J.S. implemented the protocol for ion identification. S.R. and M.A.B. carried out MD simulations. S.R., J.P.N., and G.S.S. generated mutant NIS proteins and carried out functional assays. S.R. and N.C. wrote the manuscript with input from all the other authors.

## Methods

### Cell Culture

COS-7 and 293F cell lines obtained from the American Type Culture Collection were cultured in DMEM media (GIBCO) supplemented with 10% FBS (GIBCO), 10 mM glutamine, and 100 U/mL penicillin/streptomycin (GIBCO). 293F cells in suspension were maintained in Free-Style Media (GIBCO) supplemented with 2% FBS.

### Site-Directed Mutagenesis

F67, S69, Q72, M90, Q94, Y144, V293, S416, and F417 NIS mutants were generated using as a template HA-tagged NIS cloned into pcDNA3.1. A PCR was performed with primers carrying the relevant specific codon using Kod Hot Start DNA polymerase (Novagen). After digestion with DpnI (NEB) to eliminate parental DNA, the PCR product was transformed into XL1 Blue Supercompetent *E*.*coli*. All constructs were sequenced to verify the specific nucleotide substitutions that had been made.

### Flow Cytometry

Paraformaldehyde-fixed cells were incubated with 0.1 μg/mL anti-HA (YPYDVPDYA) antibody (Roche Applied Science) in PBS/BSA (0.2%) followed by 50 nM of PE-conjugated goat anti-rat antibody (Life Technologies). The fluorescence of 50,000 cells per sample was assayed with a FACSCalibur flow cytometer (BD Bio-sciences). Data were analyzed using FlowJo software (Tree Star)^1^.

### Whole-Cell Iodide Transport

Cells were transiently transfected in 10-cm plates using 4 μg HA-tagged NIS in pcDNA3.1 and Lipofectamine/Plus reagent (Life Technologies) or 25 μg cDNAs and PEI MAX (Polyscience). After 24 hrs, they were split into 24 well/plates, and after 48 hrs they were washed twice with HBSS [140 mM NaCl, 5.4 mM KCl, 1.3 mM CaCl_2_, 0.4 mM MgSO_4_, 0.5 mM MgCl_2_, 0.4 mM Na_2_HPO_4_, 0.44 mM KH_2_PO_4_, 5.55 mM glucose, and 10 mM Hepes (pH 7.4)] following a previously published protocol^1^. For steady-state experiments, cells were incubated with HBSS containing 10, 20, 100, or 200 μM KI supplemented with carrier-free ^125^I at a specific activity of 50 μCi/μmol, at 37°C for 45 min in a CO_2_ incubator. For I^-^-dependent kinetic analysis, cells were incubated in HBSS buffer containing the I^-^ concentrations indicated (1.2-600 μM) for 2 min. Cells were then washed with HBSS, lysed with ice-cold ethanol, and the intracellular ^125^I^-^ was quantitated using a Cobra III Gamma Counter (Perkin Elmer). Kinetic parameters were determined by Michaelis-Menten linear regression after subtraction of the background recorded in cells transfected with a NIS-free plasmid.

### Protein Overexpression and Purification

293F cells were transduced with a lentiviral vector containing a rat NIS cDNA construct with an HA tag at the N-terminus and a His8 tag and a streptavidin-binding protein (IBA GmbH) tag at the C-terminus, and they were enriched for positive cells by flow cytometry via HA tag. Up to 97% of positive cells were then grown in suspension^2^.

At a density of 2.5 × 10^6^ cells/ml, 293F cells were collected and resuspended in a buffer containing 75 mM Tris⋅HCl (pH 8.0), 350 mM NaCl, and protease inhibitor cocktail (Roche), and disrupted using an Emulsiflex C3 homogenizer (Avestin). The cell debris was removed by centrifugation at 10,000 × *g* for 10 min, and the membrane-containing supernatant was collected and subjected to ultra-speed centrifugation at 250,000 × *g* for 3 hrs. Membranes were resuspended in the extraction buffer containing 75 mM Tris⋅HCl (pH 8.0), 350 mM NaCl, and protease inhibitor cocktail (Roche), and incubated with 0.5% LMNG and 0.5% GDN for 2 hrs at 4°C. NIS was subsequently purified via Strep-Tactin^®^ affinity chromatography and size exclusion chromatography via Superdex^®^ 200 Increase in the presence of 350 mM NaCl, 75 mM Tris⋅HCl (pH 8.0), and 0.005% LMNG/GDN. The fraction corresponding to the peak of the chromatogram was then used for EM preparation.

### Cryo-Sample Preparation

All cryo-EM grids were prepared by applying 2.5 μl of protein at ∼2 mg/ml to freshly glow-discharged Quantifoil 1.2/1.3, 200 mesh (Electron Microscopy Sciences), blotted for 4-6 s at 100% humidity and 7°C, and plunge-frozen in liquid nitrogen-cooled liquid ethane using a Vitrobot Mark IV (ThermoFisher Scientific).

### Data Acquisition and Processing

All the cryo-EM datasets were collected using a Titan Krios G2 electron microscope (Yale University Cryo-EM Resource) operated at 300 kV and equipped with a GIF Quantum LS imaging filter (Gatan, lnc.) and a K3 summit direct electron detector (Gatan, lnc.). Images were acquired at a superresolution pixel size of 0.534 Å. The dose rate was 16.1 e/physical pixel/s, yielding a total dose of 45 e/Å^2^. Movie stacks were drift-corrected using MotionCorr2^3^ with a 5 × 5 patch and twofold binning. Contrast transfer function (CTF) information was estimated from dose-weighted images by CTFFind4.1^4^.

Two datasets for NIS in the presence of its substrate Na^+^ were acquired in two separate sessions from two different protein preparations, yielding 3861 and 4164 movies respectively. Each dataset was analyzed individually. Movie stacks were drift-corrected using MotionCorr2^3^ with a 5 × 5 patch and a twofold binning. Contrast transfer function (CTF) information was estimated from dose-weighted images by CTFFind4.1^4^ in Relion^5^ with exhaustive searching. Particles were picked by Relion Auto-picking using Laplacian-of-Gaussian blob detection. After particles were extracted at a pixel size of 2.73 Å/px, 3 rounds of 2D classification were performed using the fast subset for a larger dataset followed by one with all data. After the good classes were selected, the contributing particles were re-extracted at the original size (1.068 Å/px), and multiple rounds of 3D classification and 3D auto-refinement were performed in Relion. The final particles leading to the best 3D-refined maps in both datasets were combined and subjected to additional 3D classification and 3D Refinement. The final pool of 108k selected particles was extracted with different particle-box sizes and imported in cryoSPARC^6^. Non-Uniform Refinement^7^ of the several particle sets led to the choice of the optimal particle-box size (288 px)^8^; subsequent Global CTF Refinement and Non-Uniform Refinement resulted in a 3.47 Å map. The NIS model was built in Coot^9^ and refined in Phenix^10^ Real Space Refine. Local Resolution analysis was performed in cryoSPARC. We used Chimera^11^ to generate a mask around one of the monomers and cryoSPARC to locally refine the map, thereby obtaining a final 3.29 Å map. The model was then further refined in Phenix, yielding the final structure.

A dataset for NIS in the presence of both its substrates Na^+^ and I^-^ (purification buffer supplemented with 2 mM KI) was acquired for a total of 3867 movies. Particles were chosen by Relion Auto-picking using Laplacian-of-Gaussian blob detection, for a total of 3 × 10^6^ particles followed by 3 rounds of 2D classification using the fast subset for a larger dataset and then one last one with all data. After class selection, the particles were re-extracted at the original size (1.068 Å/px), and multiple rounds of 3D classification and 3D auto-refine were performed in Relion 3.0.8. The final pool of particles was then re-extracted with a different box size and imported in cryoSPARC^6^. Non-Uniform Refinement^7^ of the several particle sets led to the choice of the optimal particle-box size (288 px)^8^; subsequent Global CTF Refinement and Non-Uniform Refinement resulted in a 3.46 Å map. The NIS model was built in Coot^9^ and refined in Phenix^10^ Real Space Refine. Local Resolution analysis was performed in cryoSPARC using Local Resolution Estimation, and the density maps were prepared using UCSF Chimera^11^. We used Chimera to generate a mask around one of the monomers and cryoSPARC to locally refine the map, thereby obtaining a final 3.12 Å map. The model was then further refined in Phenix, yielding the final structure.

A dataset for NIS in the presence of both its substrates Na^+^ and ReO_4_^-^ (purification buffer supplemented with 1 mM NaReO_4_) was acquired, for a total of 4431 movies. Particles were chosen by Relion Auto-picking using Laplacian-of-Gaussian blob detection, for a total of 3 × 10^6^ particles, followed by 3 rounds of 2D classification using the fast subset for a larger dataset and then one last one with all data. After class selection, the particles were re-extracted at the original size (1.068 Å/px), with a different box size and imported in cryoSPARC^6^. Several rounds of Heterogeneous Refinement lead to a final 432k particles that were then subjected to Non-Uniform Refinement^7^ and subsequent Global CTF Refinement to obtain a 3.24 Å map. The NIS model was built in Coot^9^ and refined in Phenix^10^ Real Space Refine. Local Resolution analysis was performed in cryoSPARC. Symmetry expansion and local refinement applying a mask covering a single monomer led to a map with 3.15 Å resolution. The NIS model was then refined in Phenix^10^ Real Space Refine.

### Ion identification

As a cryo-EM density map reflects electrostatic potentials, we sought to quantitate the density in the vicinity of the ion sites. We ascertained the I^-^ and ReO_4_^-^ map densities along 24 lines (48 equally spaced directions generated using HealPix pixelation^12^) centered on each ion position. Densities along the lines were evaluated by trilinear interpolation on a four-times Fourier-upsampled density map. An overall spherically averaged density (as a function of radial distance from the ion) was also computed from the upsampled map. To aid comparison of numerical values between maps, density values in each map were normalized using the peak density at the α-carbon positions of the residues Q72 and Q94^12,13^.

### Molecular Dynamics Simulations

Molecular dynamics simulations were carried out using the NIS apo- and holo-structures (2 Na^+^/I^-^ and Na^+^ structures) embedded in a pre-equilibrated 1,2-Dioleoyl-sn-glycero-3-phosphocholine (DOPC) bilayer. We obtained the parameters and scripts for the minimization, equilibration, and production in GROMACS using CHARMM-GUI^14^. The composition of each system in the production runs is summarized in Supplemental Information Table SX. All simulations were carried out with GROMACS, version 2020^15^, in conjunction with the CharMM36 force field^16^, using a TIP3P water model^17^, Berger-derived DOPC lipids^18^, and ion parameters by Joung and Cheatham^19^. Van der Waals interactions were cut off at 1 nm, and electrostatics were treated by PME^20^ beyond 1 nm. Temperature and pressure were kept at 310.5 K and 1 bar using the V-Rescale thermostat^21^ and Parrinello-Rahman barostat^22^, respectively. All bonds were restrained using LINCS^23^, and an integration time step of 2 fs was used. We energy-minimized (steepest descent) using a double-precision version of GROMACS, and six steps (125, 125, 125, 500, 500, and 500 ns) of position restraint with a gradually lifted harmonic force constant (Fc) with different values for backbone atoms (100000, 2000, 1000, 500, 200, and 50 kJ/mol/nm^2^), side-chain atoms (2000, 2000, 1000, 500, 200, 50, and 0 kJ/mol/nm^2^), residue dihedrals (1000, 200, 200, 100, and kJ/mol/nm^2^), and lipids (1000, 400, 400, 200, 40, and 0 kJ/mol/nm^2^). Van der Waals parameters for I^-^ were taken from Li et al^24^. Trajectories and the Jupyter notebook used for the calculations are available upon request.

---

**Extended Data Fig. 1.**
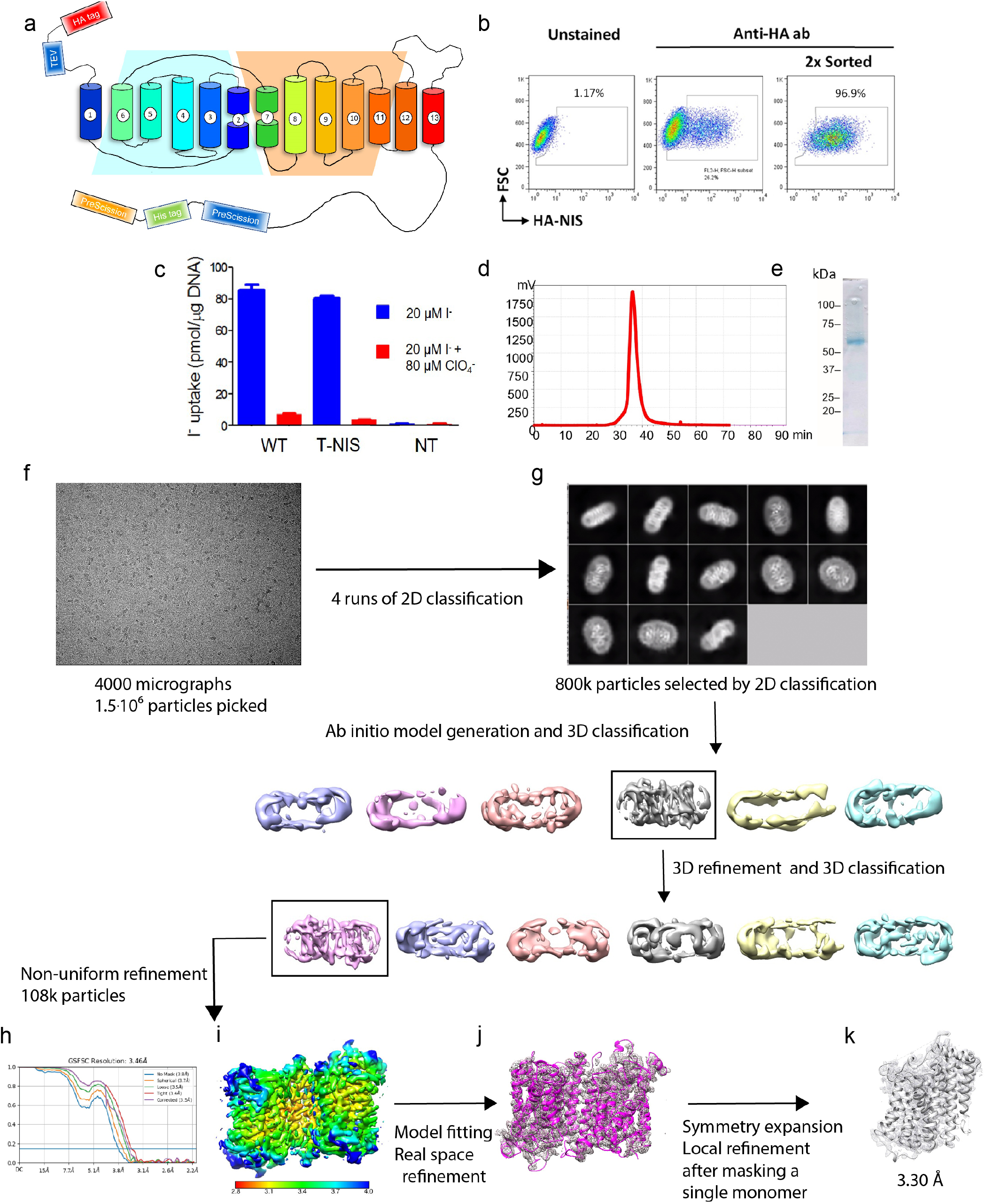
Cryo-EM data processing and determination of the structure of NIS. Topology model of the engineered NIS molecule (rt-NIS) whose cDNA was used to transduce 293F cells. We engineered an HA tag onto the N-terminus and a HIS and an SBP (streptavidin-binding protein) tag onto the C-terminus for affinity purification. The tags are separated from NIS by a TEV protease site at the N-terminus and a PreScission site at the C-terminus. **b**. Enrichment of 293F cells expressing NIS by flow cytometry using an anti-HA antibody. After two rounds of sorting, >96% of the cells expressed NIS at the plasma membrane. **c**. I^-^ transport assay. rtNIS transports virtually as much I^-^ as WT NIS does, whereas nontransduced (NT) cells transport no I^-^**d**. Size exclusion chromatography (SEC) of the fraction of NIS purified in LMNG/GDN used for cryo-EM imaging detected by Trp-fluorescence. **e**. Coomassie blue staining of SEC-purified NIS subjected to SDS-PAGE. **f**. Cryo-EM micrograph of apo-NIS (representative of 8025 micrographs that yielded similar results). **g**. Selected 2D class averages obtained after 4 rounds of 2D classification using Relion (top image) and data-processing workflow (bottom two rows). **h**. NIS dimer map colored according to local resolution. **i**. Fitting of NIS sequence to the electron density map. **j**. Local refinement. **k**. Fourier shell correlation (FSC) of the locally refined map.

**Extended Data Fig. 2.**
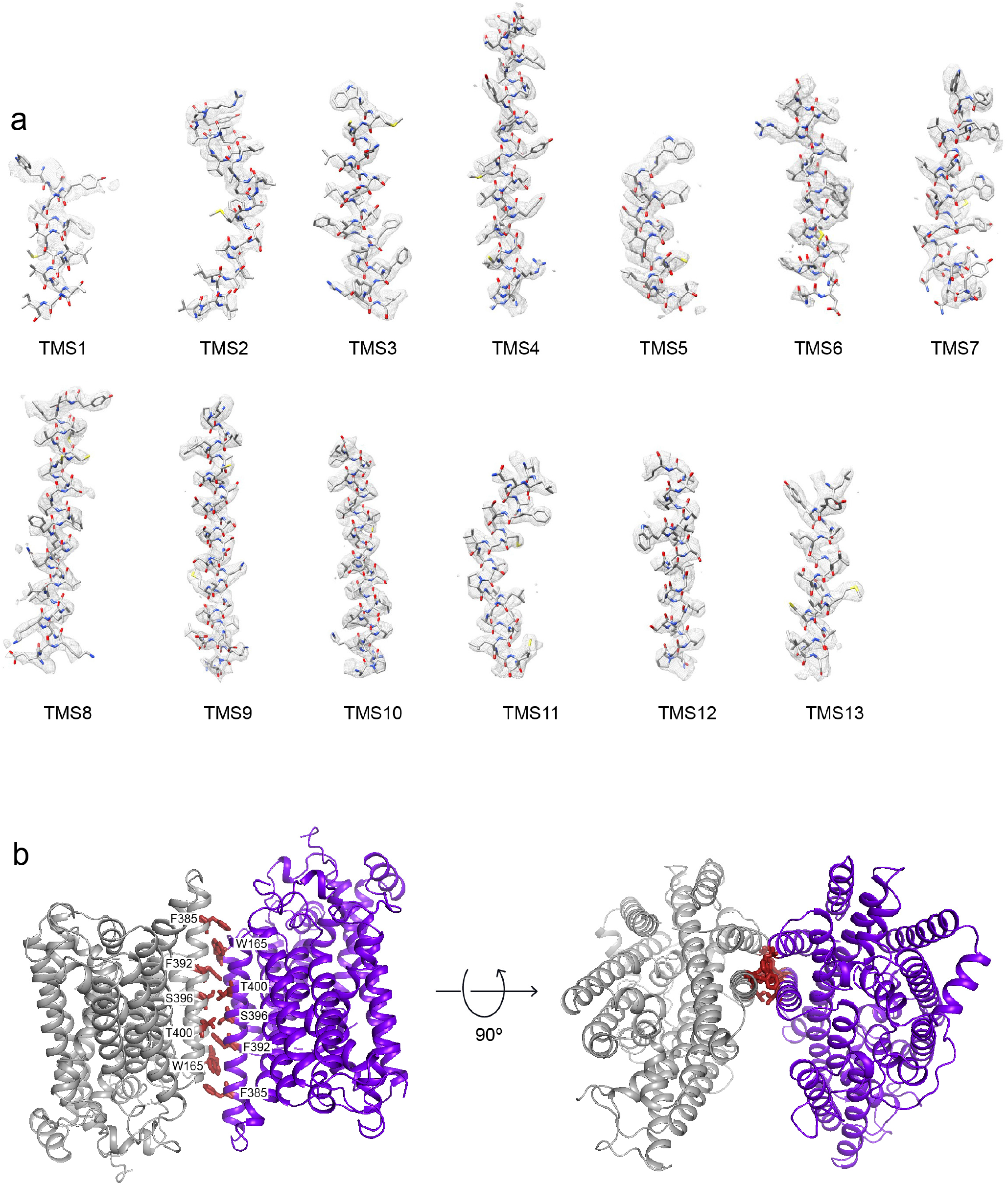
Cryo-EM densities and model of TMSs and dimer assembly of NIS. **a**. α-helical features are clearly visible in all 13 TMSs. **b**. Residues interacting at the dimer interface in apo-NIS.

**Extended Data Fig. 3.**
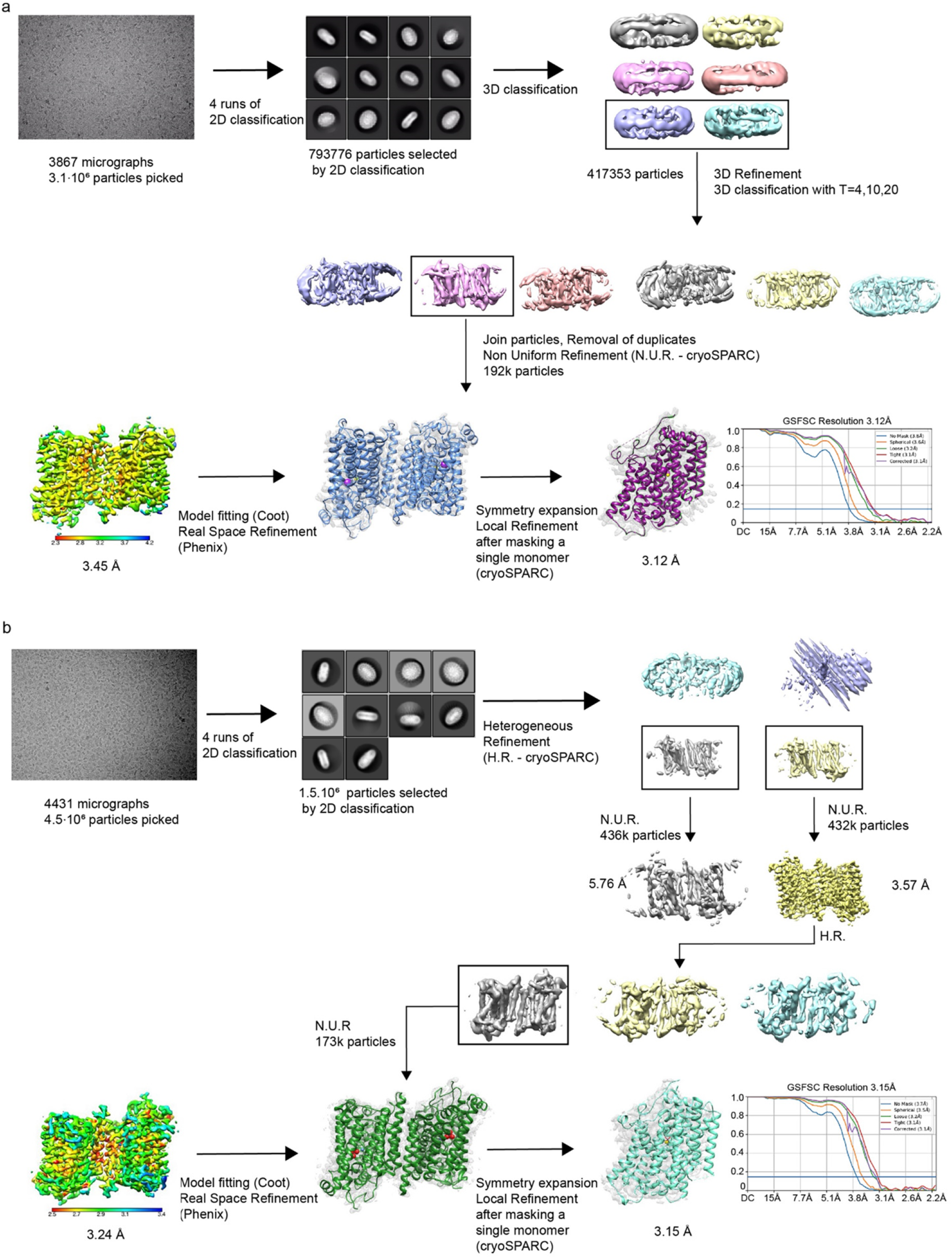
Cryo-EM data processing and determination of the structures of NIS with substrates bound. **a**. NIS-I^-^. **b**. NIS-ReO_4_^-^.

**Extended Data Fig. 4.**
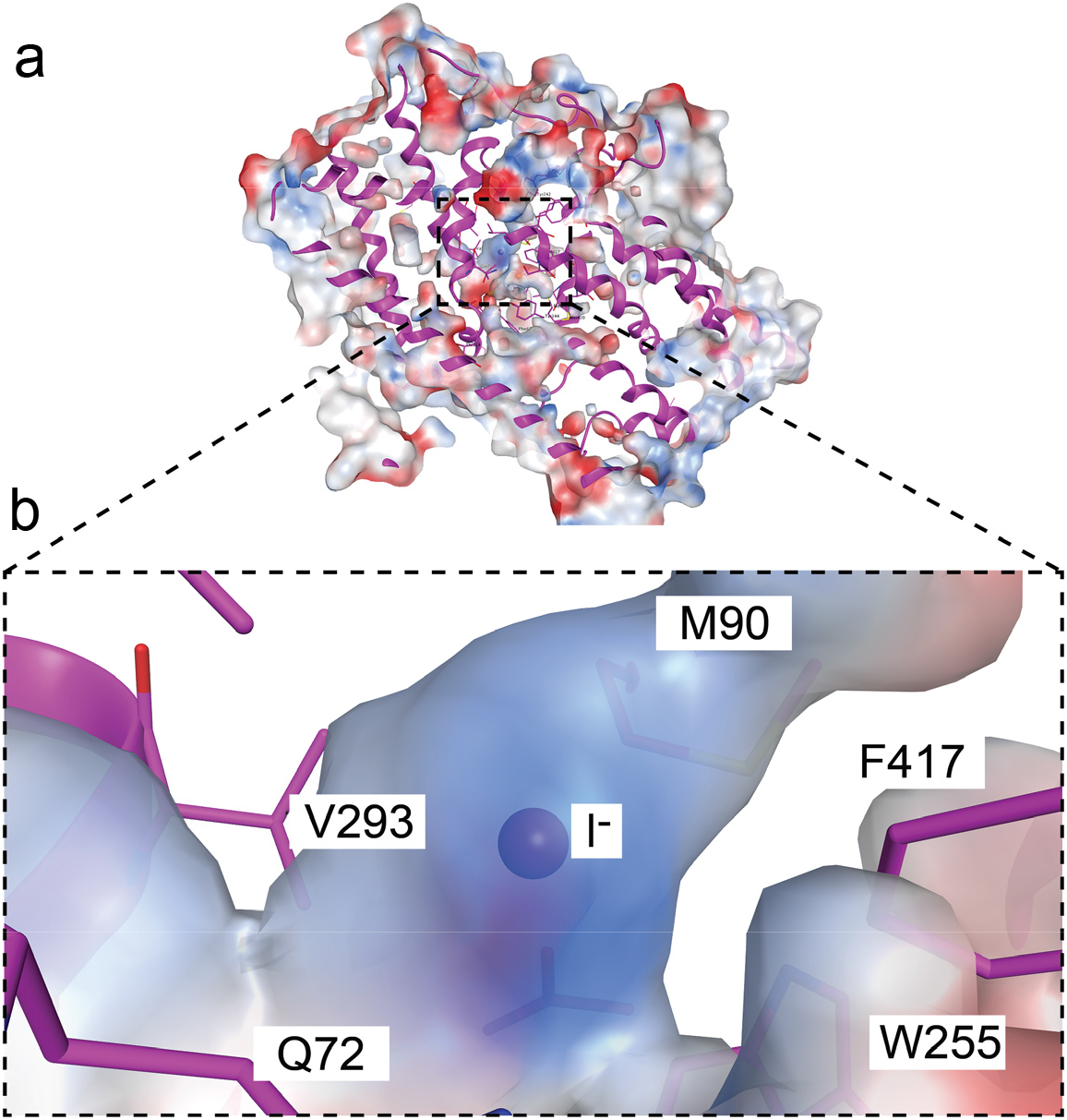
I^-^ binds to a partially positively charged cavity. **a**. NIS-I^-^ side view. The solvent accessible surface is colored according to the electrostatic potential in k_B_T/e (red, negative; blue, positive) and cropped to expose the I^-^ binding cavity. **b**. Close-up of the I^-^ binding cavity showing the positive nature of its surface.

**Extended Data Fig. 5.**
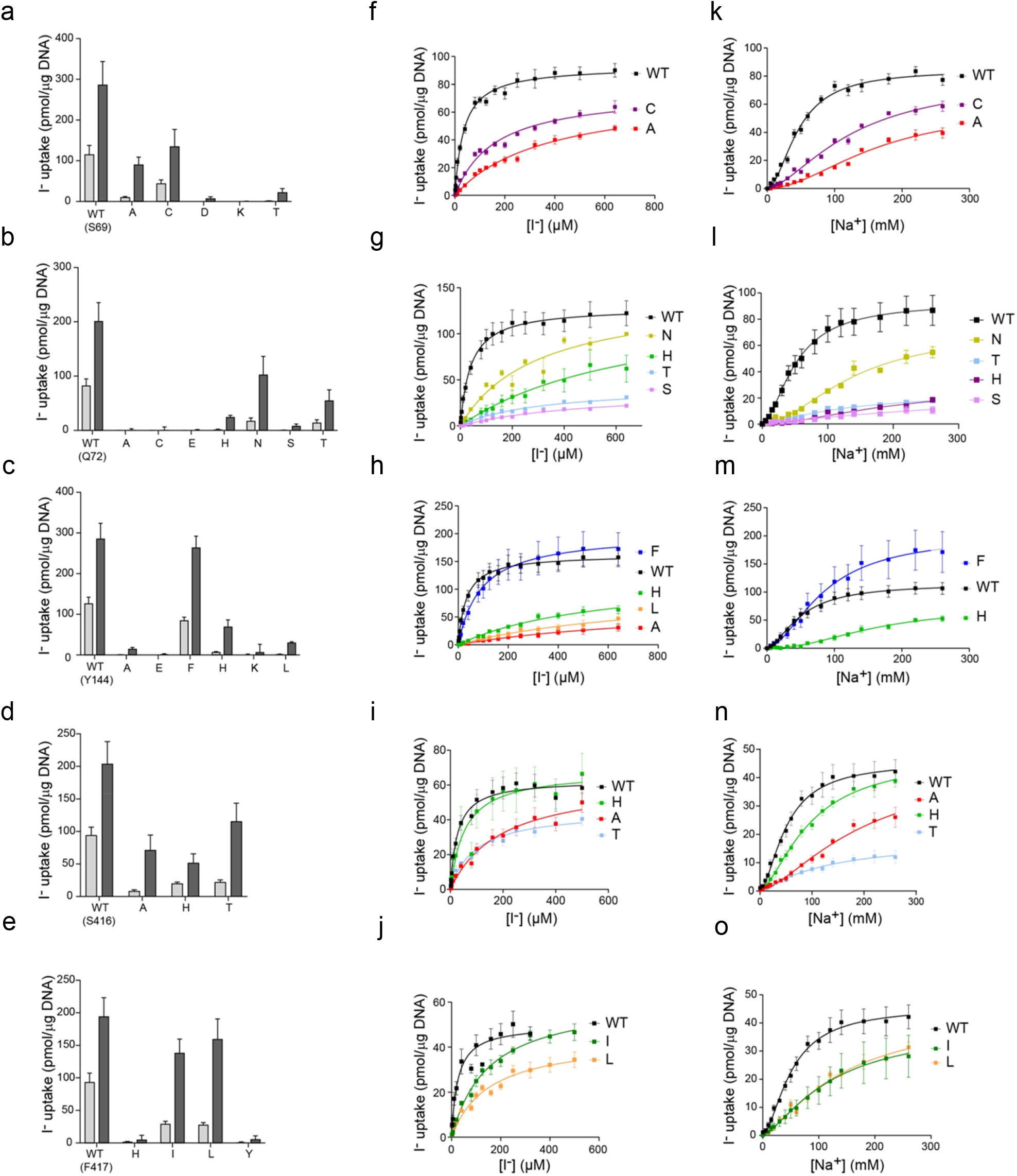
Effects of single amino acid substitutions at positions 69, 72, 144, 416, 417 on iodide transport. **a-e**. NIS-mediated I^-^ uptake at steady state. cDNA constructs coding for NIS mutants were transfected into COS7 or HEK cells. I^-^ uptake by these NIS mutants was measured at 20 and 200 μM I^−^ at 140 mM Na^+^ for 30 min with or without the NIS-specific inhibitor ClO_4_^-^ to determine NIS-mediated transport (values obtained in the presence of ClO_4_^-^, which are < 10% of the values obtained in its absence, have already been subtracted). Results are expressed as pmol of I^-^ accumulated/μg DNA ± SE. Values represent averages of the results from two or three different experiments, each of which was carried out in triplicate. **f-j**. Kinetic analysis of initial rates of I^-^ uptake (2-min time points) determined at 140 mM Na^+^ and varying concentrations of I^-^. **k-o**. Kinetic analysis of initial rates of I^-^ uptake (2-min time points) determined at varying concentrations of extracellular Na^+^.

**Extended Data Fig. 6.**
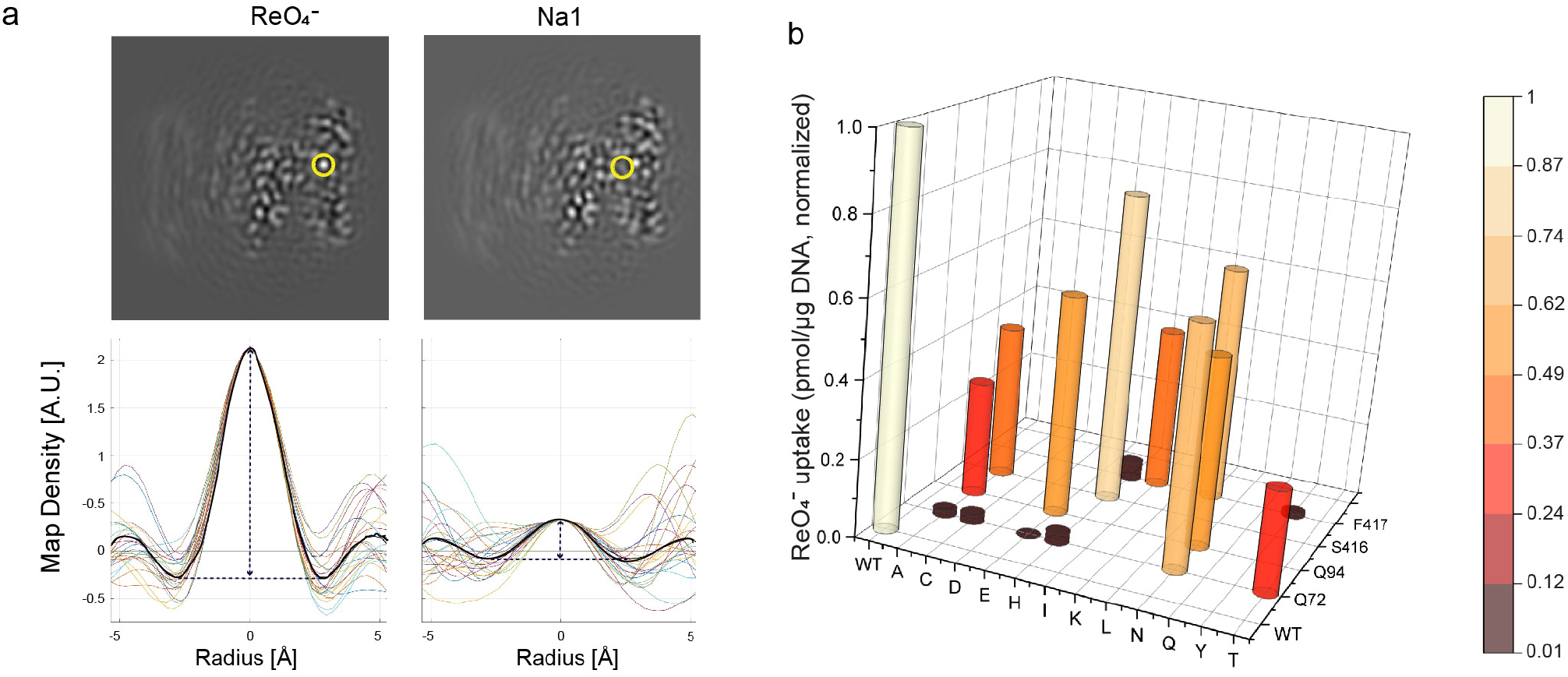
Identification of the ions in the NIS-ReO_4_^-^ structure. **a**. The ions transported by NIS were identified by evaluating the map density along 24 lines passing through each site (circled in yellow on images of map slices). Values are plotted for each line and the spherically-averaged mean is plotted in black (lower panels). **b**. Effects of substitutions in binding-site residues on ReO_4_^-^ transport at steady state (measured at 3 μM ReO_4_^-^ and 140 mM Na^+^ for 30 min (values obtained in the presence of ClO_4_^-^, which are < 10% of the values obtained in its absence, have already been subtracted). Values are normalized to those obtained with WT NIS.

**Extended Data Fig. 7.**
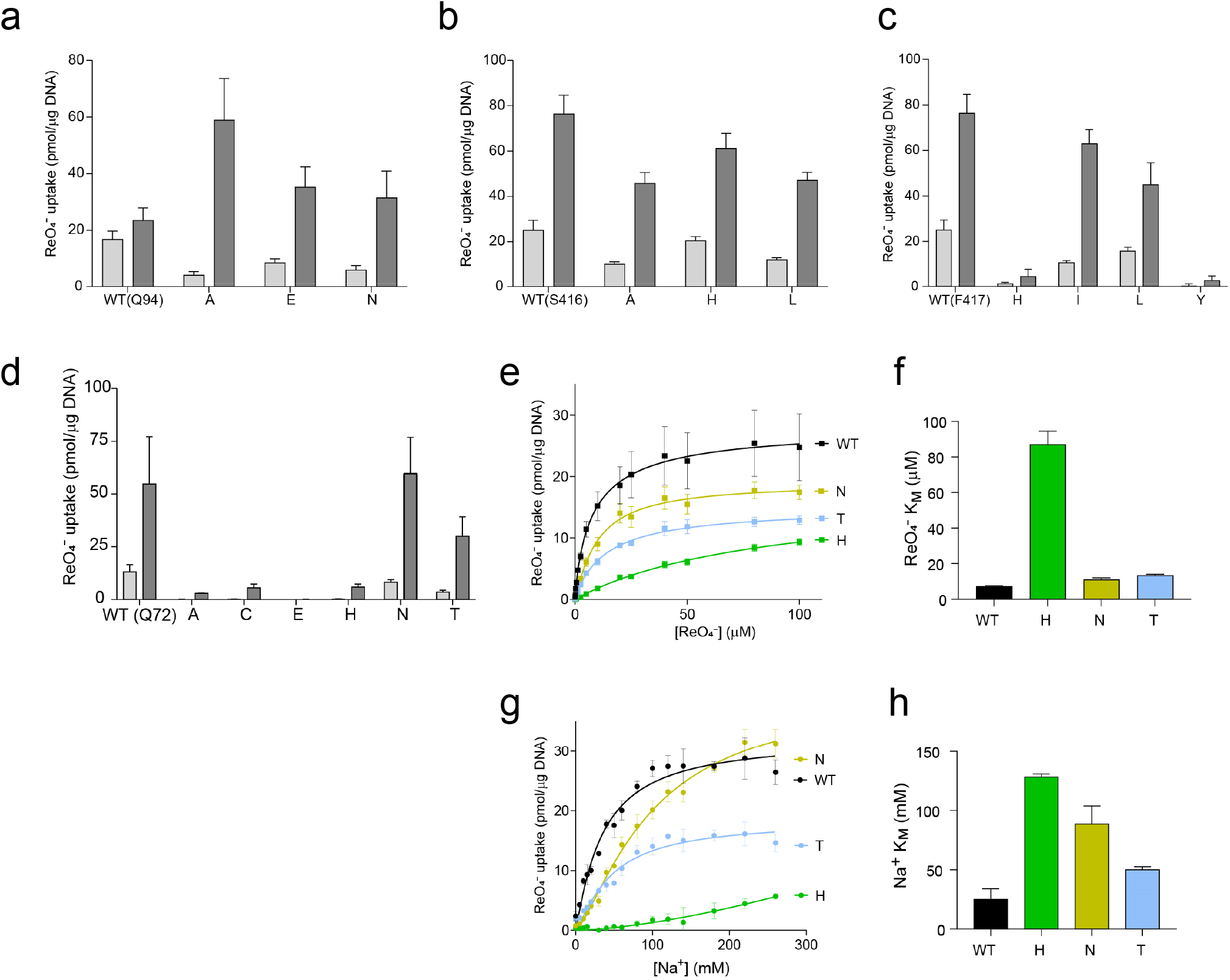
Effects of single amino acid substitutions at positions 72, 94, 416 and 417 on ReO_4_^-^ transport. **a-d**. NIS-mediated ReO_4_^-^ uptake at steady state. cDNA constructs coding for NIS mutants in which Q72 is replaced with the residues indicated were transfected into COS7 or HEK cells. ReO_4_^-^ uptake by these NIS mutants was measured at 3 and 30 μM ReO_4_^-^ at 140 mM Na^+^ for 30 min with or without the NIS-specific inhibitor ClO_4_^-^ (values obtained in the presence of ClO_4_^-^ already subtracted). Results are given as pmols of ReO_4_^-^ accumulated/μg DNA ± SE. Values represent averages of the results from two or three different experiments, each of which was carried out in triplicate. **e**. Kinetic analysis of initial rates of ReO_4_^-^ uptake (2-min time points) for Q72 NIS mutants determined at varying concentrations of extracellular ReO_4_^-^ and varying concentrations of extracellular Na^+^. **f**. ReO_4_^-^ *K*_*M*_ values determined from (e). **g**. Kinetic analysis of initial rates of ReO_4_^-^ uptake (2-min time points) for Q72 NIS mutants determined at 100 μM ReO_4_^-^ and varying concentrations of extracellular Na^+^. **h**. Na^+^ *K*_*M*_values determined from (g).

**Extended Data Fig. 8.**
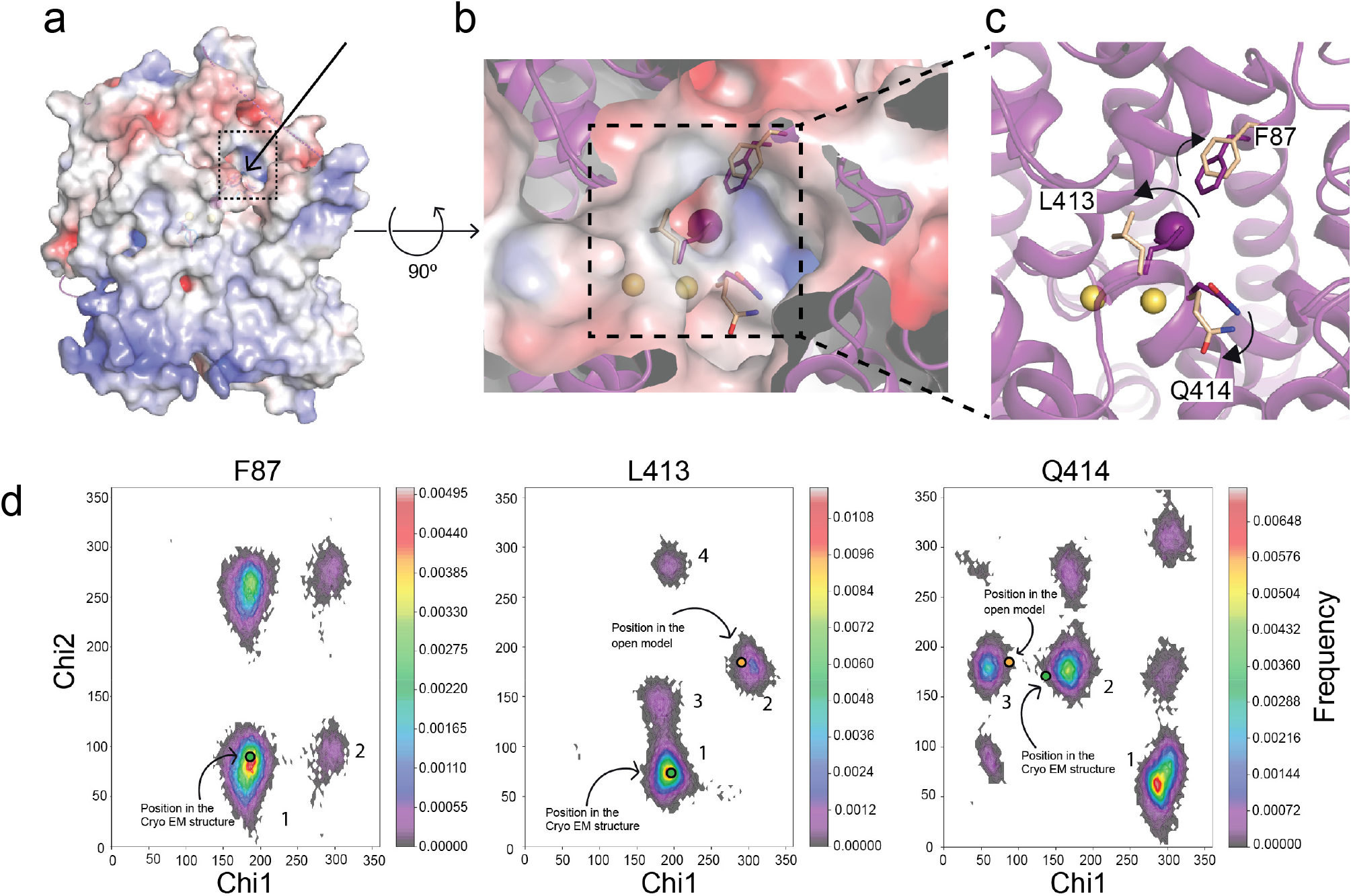
Entry pathway for the NIS substrates. **a**. Surface representation of a side view of the NIS-I^-^ structure: the arrow and the dotted square indicate the position of the proposed entry pathway. **b**. Close-up of the top view of the surface showing the substrates (Na^+^ ions represented by yellow spheres, I^-^ by a magenta sphere), and the positions of F87, L413, and Q414 in the NIS-I^-^ structure (magenta) and the models generated from MD simulations corresponding to the opening (wheat) of the substrate-binding cavity to the extracellular milieu. **c**. Magnification of the top of the substrate-binding cavity. The arrows indicate how the amino acids move away from their original positions as the cavity transitions from closed to open. **d**. Ramachandran plots of the chi-1 and chi-2 side chain dihedral angles of F87, L413, and Q414 visited during the MD simulations with NIS-I^-^. The dihedral angles selected are the principal determinants of the position of the side chain. The excursions of these dihedral angles (during the MD simulations) away from the conformational basins corresponding to the cryoEM structure (green dots in basins 1, 1, and 2, in F87, L413, and Q414, respectively) and toward conformational basins (blue dots in basins 1, 2, and 3, in F87, L413, and Q414, respectively) open up the entry path (b). In these histograms, the frequency of a given conformational state is indicated by a rainbow gradient from deep purple (0 frequency) to red (highest frequency); the most populated conformational basins are numbered in descending order of population.

**Extended Data Fig. 9.**
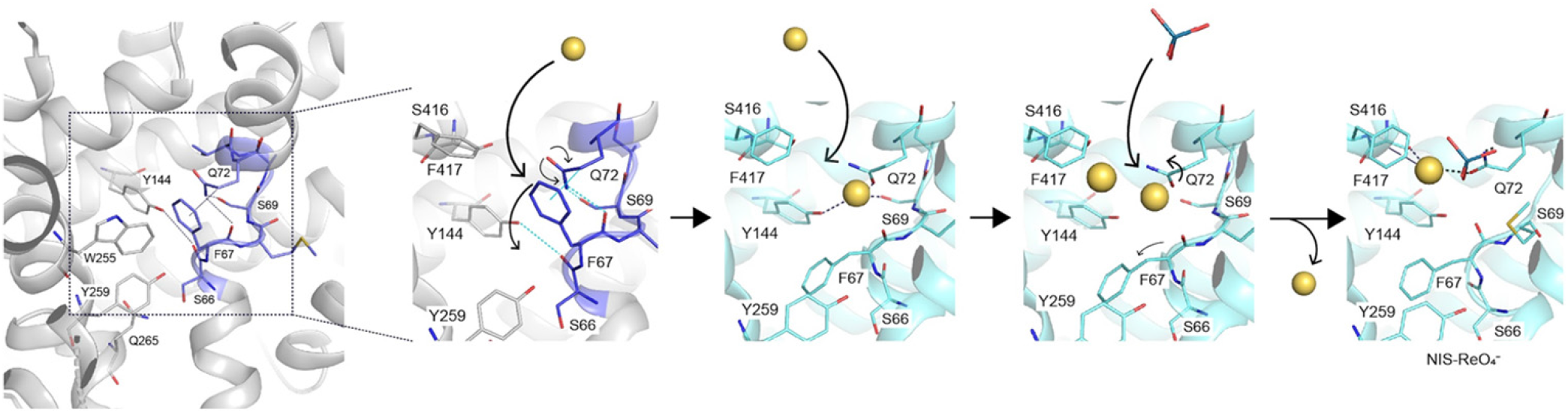
Perrhenate binding mechanism starting from the apo-NIS structure. A hydrogen-bond network between F67, S69, Q72, and Y144 and a hydrophobic stacking interaction between F67 and Q72 are disrupted by the binding of the first Na^+^. This facilitates the binding of a second Na^+^ and ReO_4_^-^, which causes the release of one Na^+^.

**Extended Data Fig. 10.**
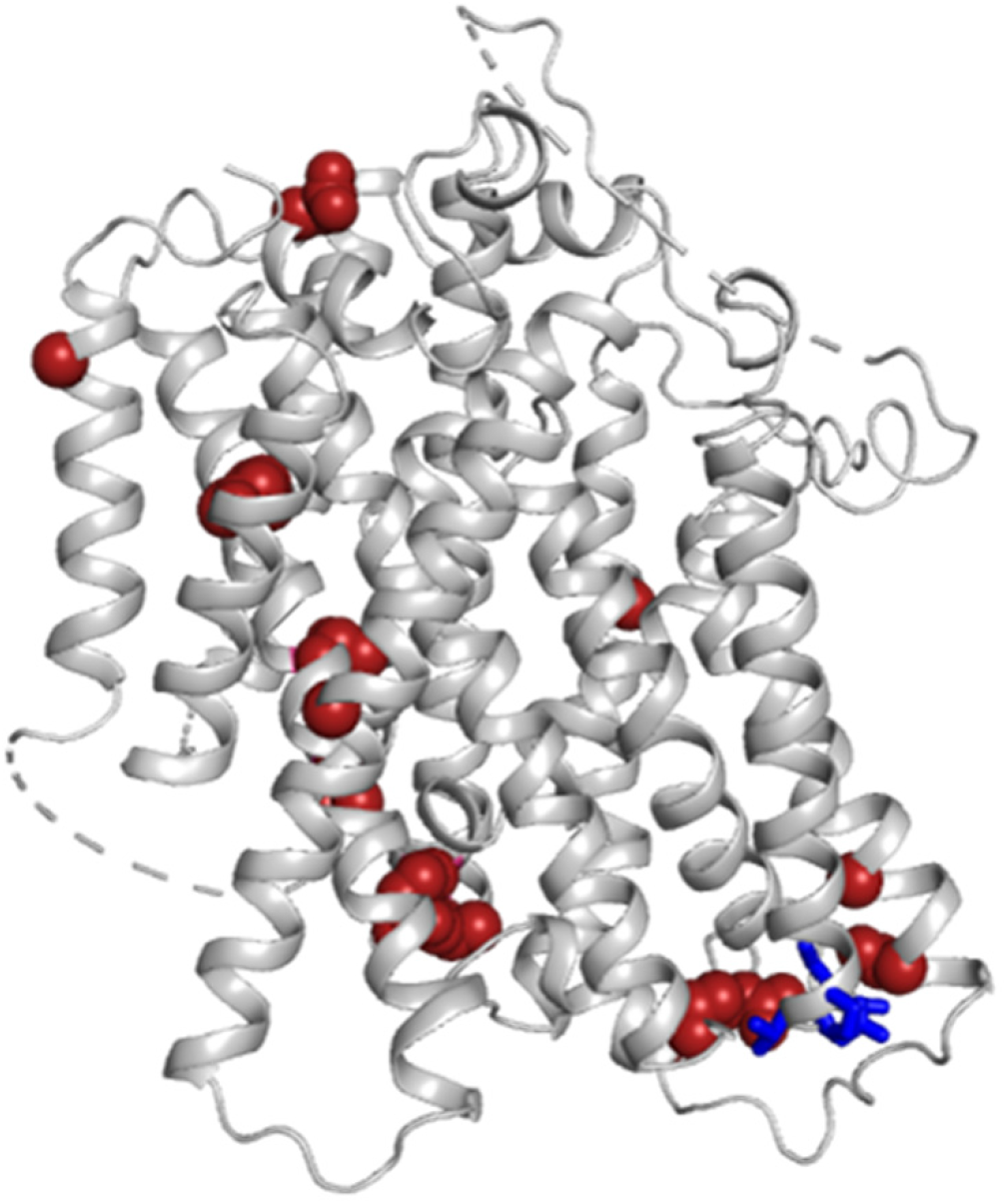
Apo-NIS structure with amino acids mutated in patients with ITDs shown as spheres. Single amino acid substitutions are indicated by red spheres; deleted residues are in blue.

**Extended Data Table 1.**
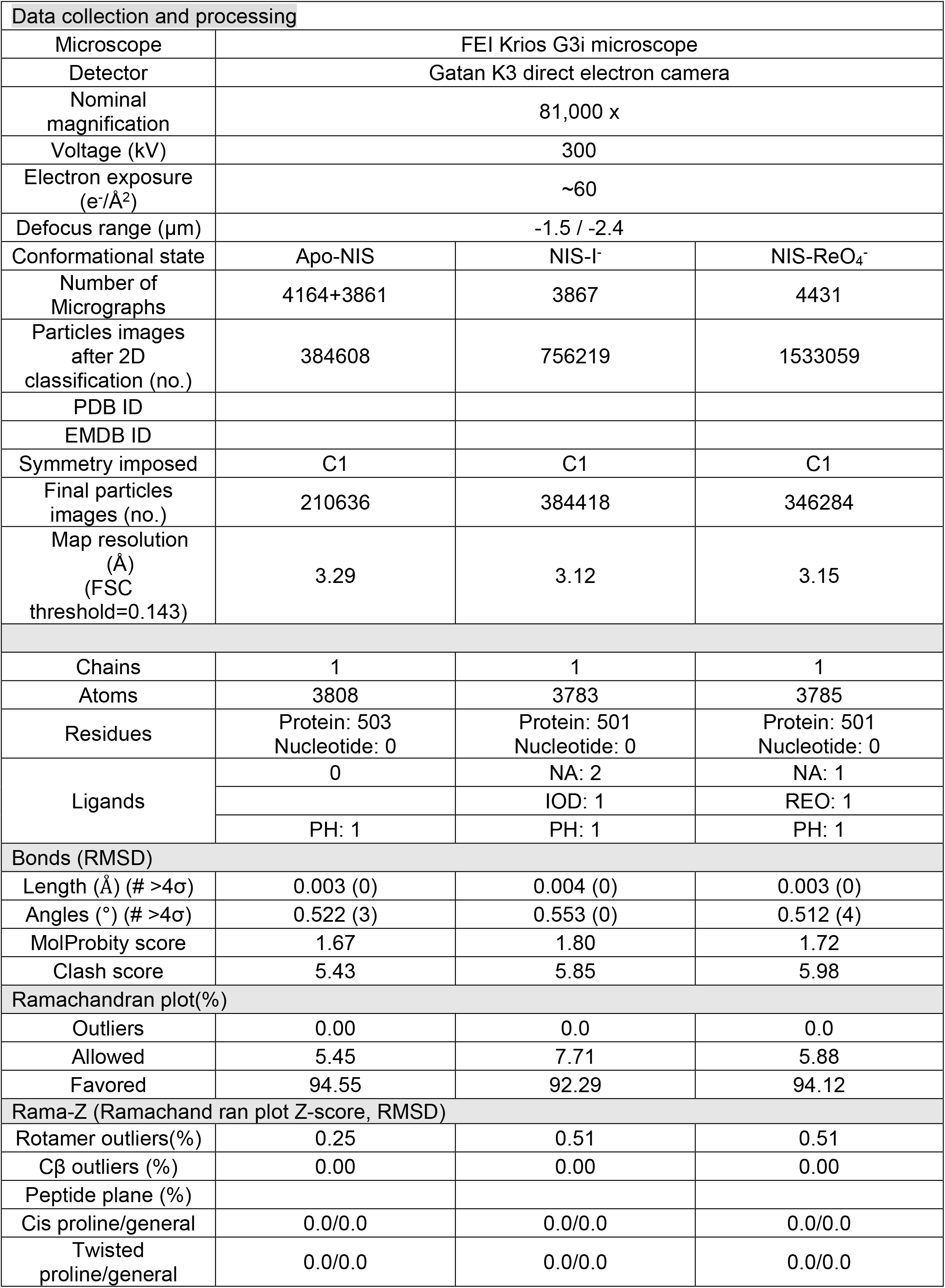
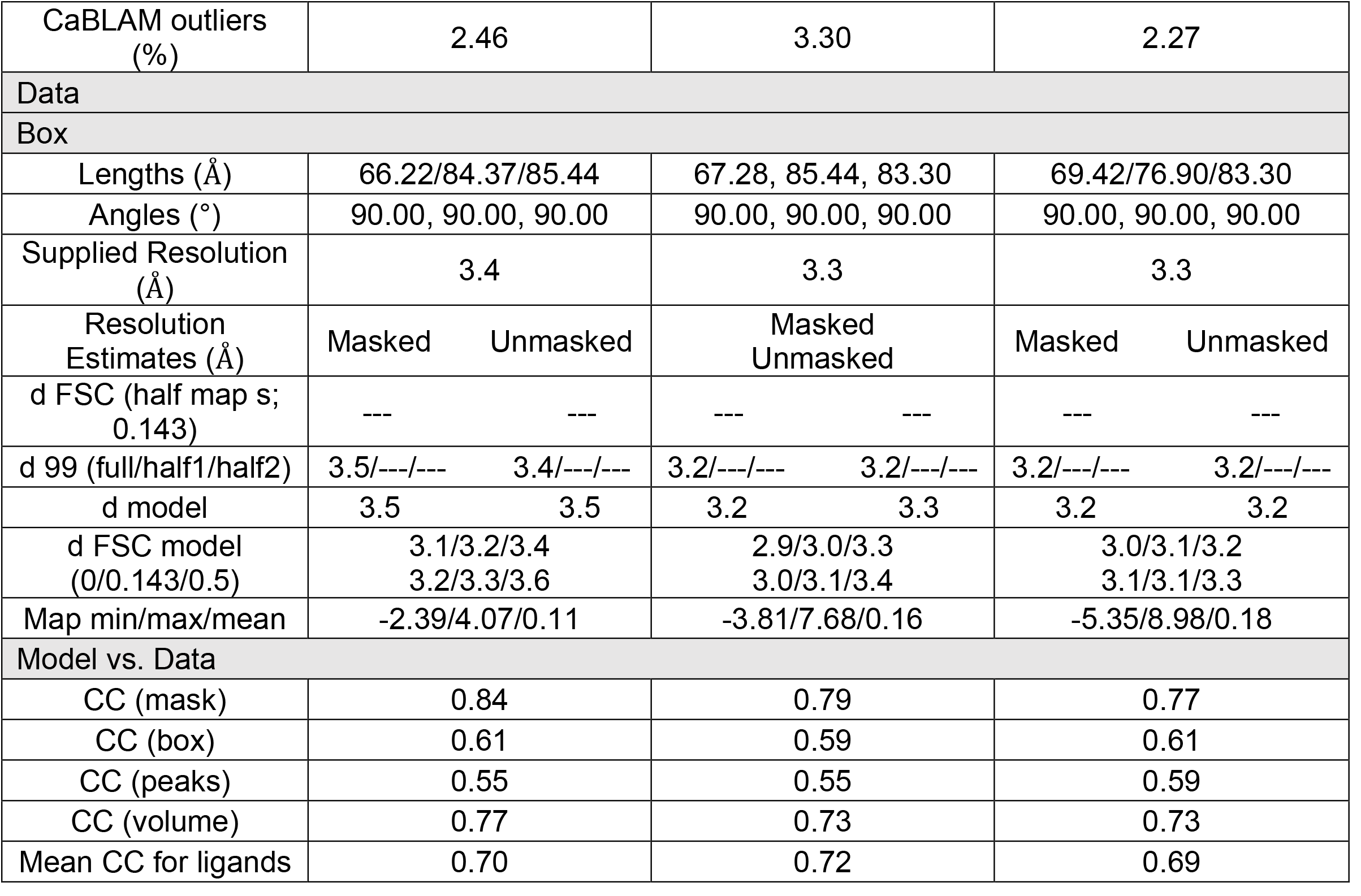
Cryo-EM data collection, refinement, and validation statistics.

