## Supplemental material for "Structural insights into the mechanism of the sodium/iodide symporter (NIS)"

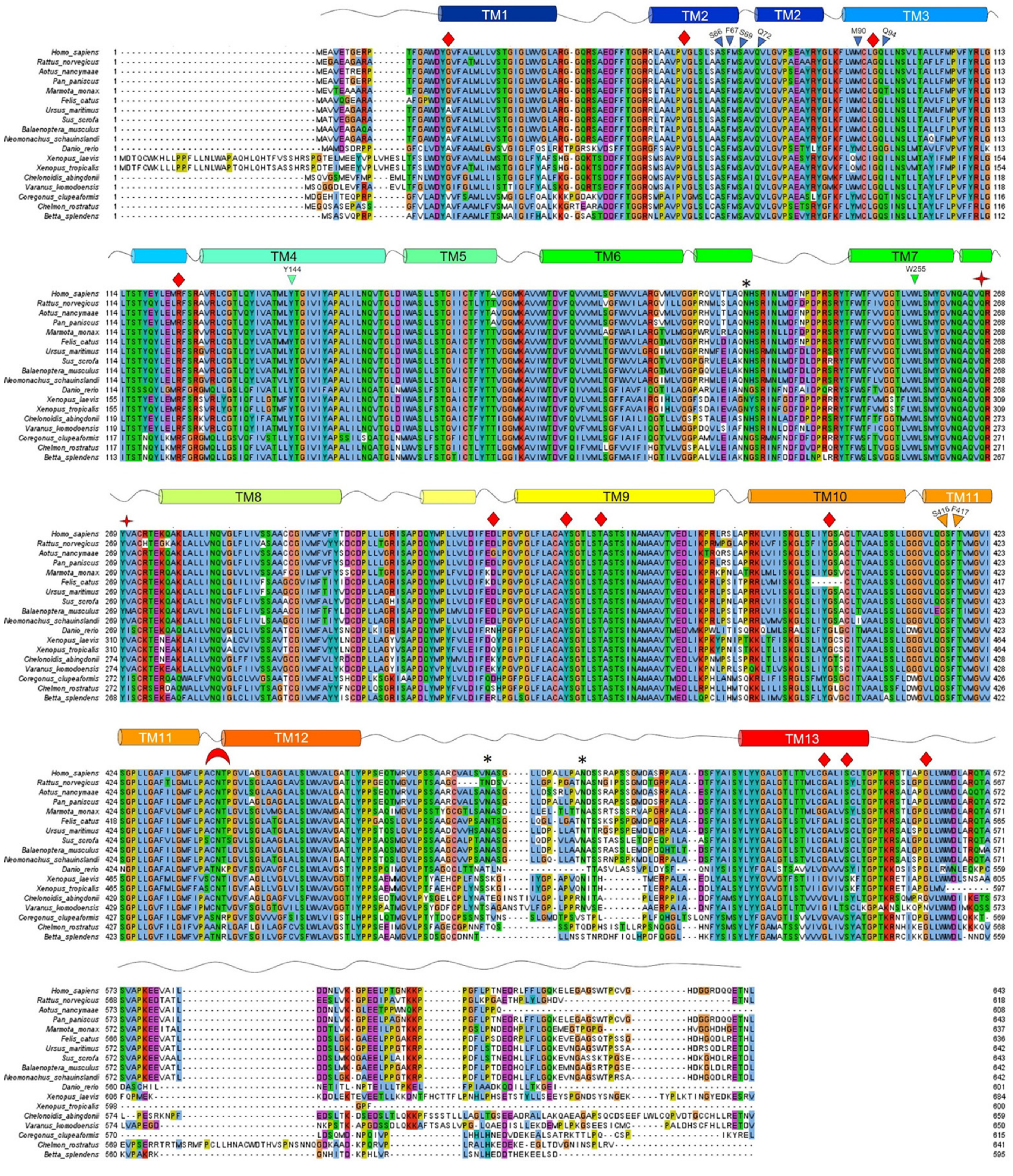


**Supplementary Fig. 1 Alignment of NIS sequences from different species.** α-helices are represented by cylinders. Glycosylation sites are marked with a *; critical residues with inverted triangles; non-WT residues found in IDD patients with diamonds; and the deletion Δ439-443 found in IDD patients with a crescent moon.

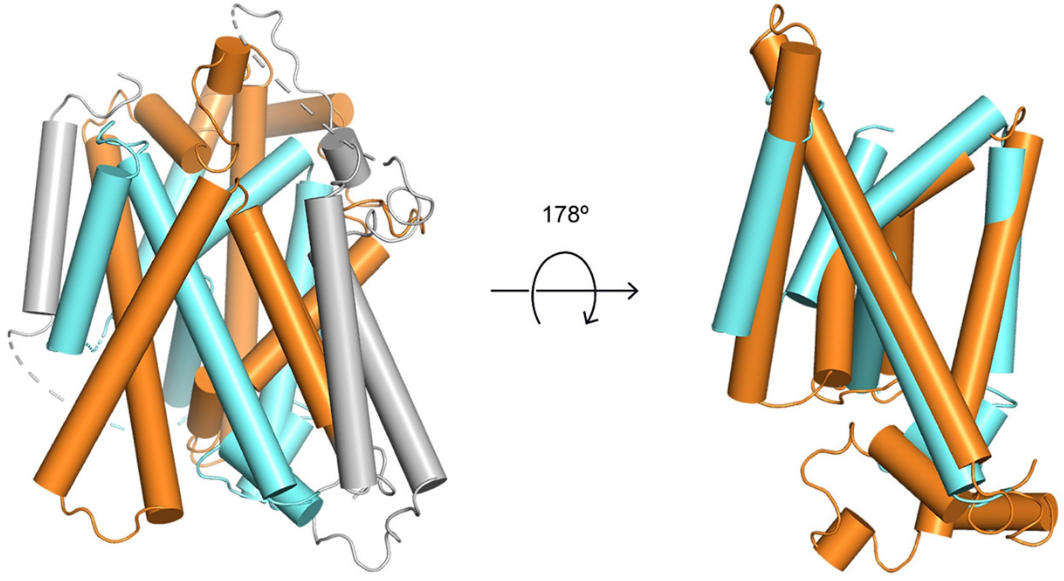


**Supplementary Fig. 2 |Alignment of the inverted structural repeats.** NIS has a LeuT fold with two 5-helix bundle domains (TMSs 2-6 and 7-11) related by a pseudo-two-fold symmetry. The alignment between the two repeats shows a root mean square deviation (RMSD) of 3.0 Å.

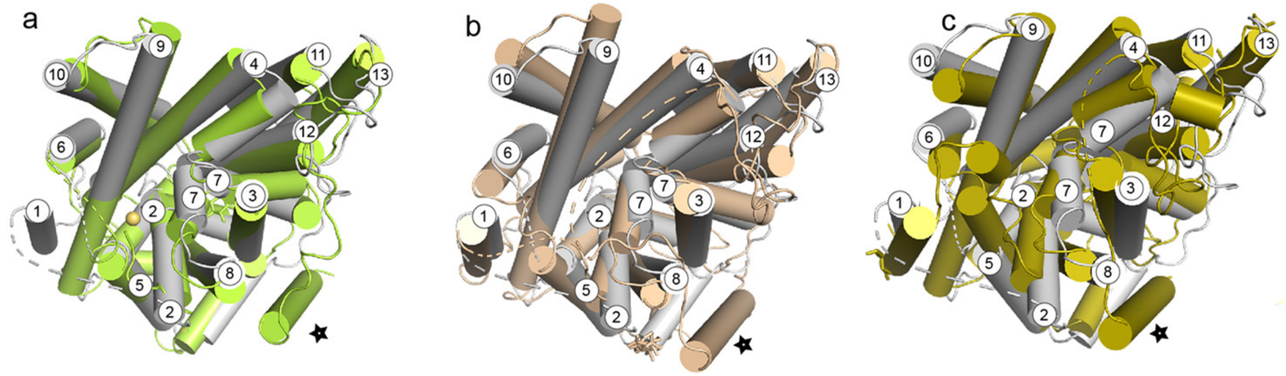


**Supplementary Fig. 3 | Structural alignment** between NIS in grey, vSGLT in green (a), hSGLT1 in wheat (b), and hSGLT2 in olive (c). *the stars in g, h, and i indicate the extra TMSs present in vSGLT, SGLT1, and SGLT2.

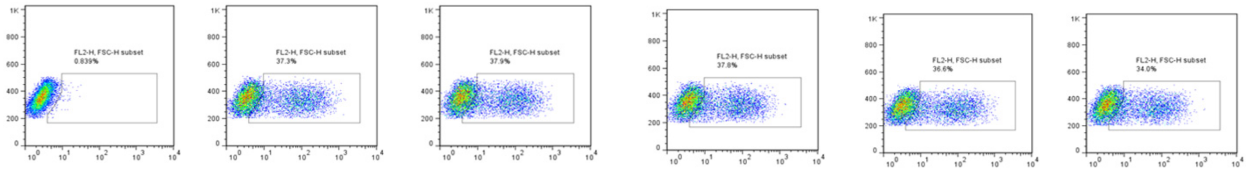

Empty vector | WT (F67) | F6 7A | F67 Q | F6 7L | F6 7E

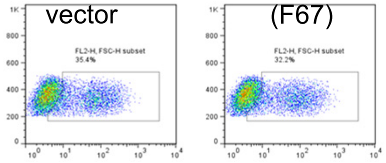

F6 7K | F67 H

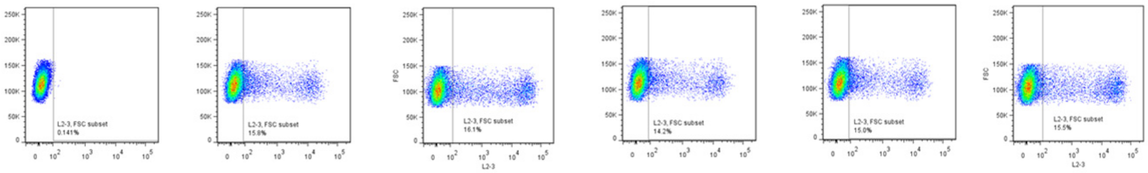

Empty | WT | F41 | F41 | F41 | F41

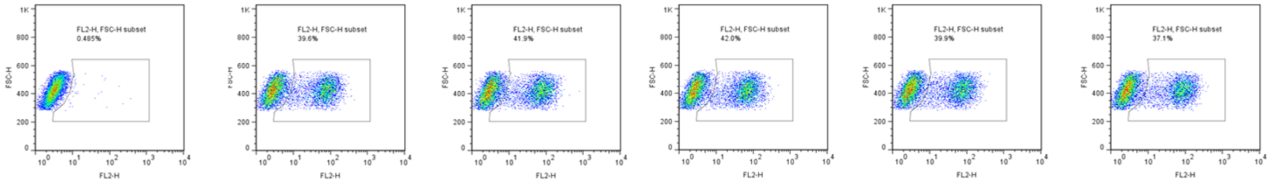

Empty | WT | Q7 2C | Q7 2H | Q7 2N | Q7 2S

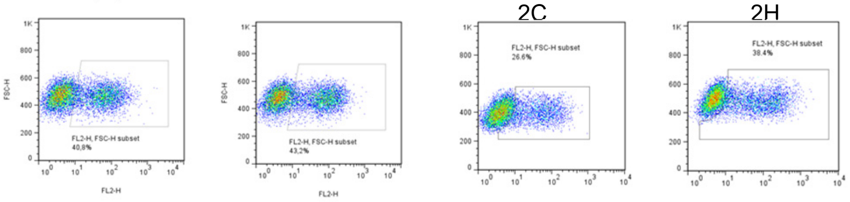

Q7 2A | Q7 2E | Q7 2T | Q7 2S

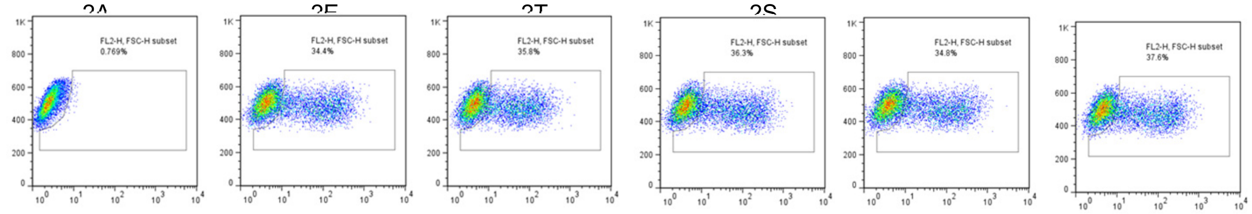

Empty vector | WT (S69) | S6 9A | S69 D | S69 C | S6 9K

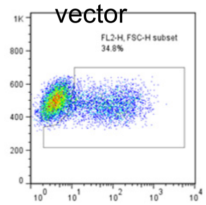

S6 9T

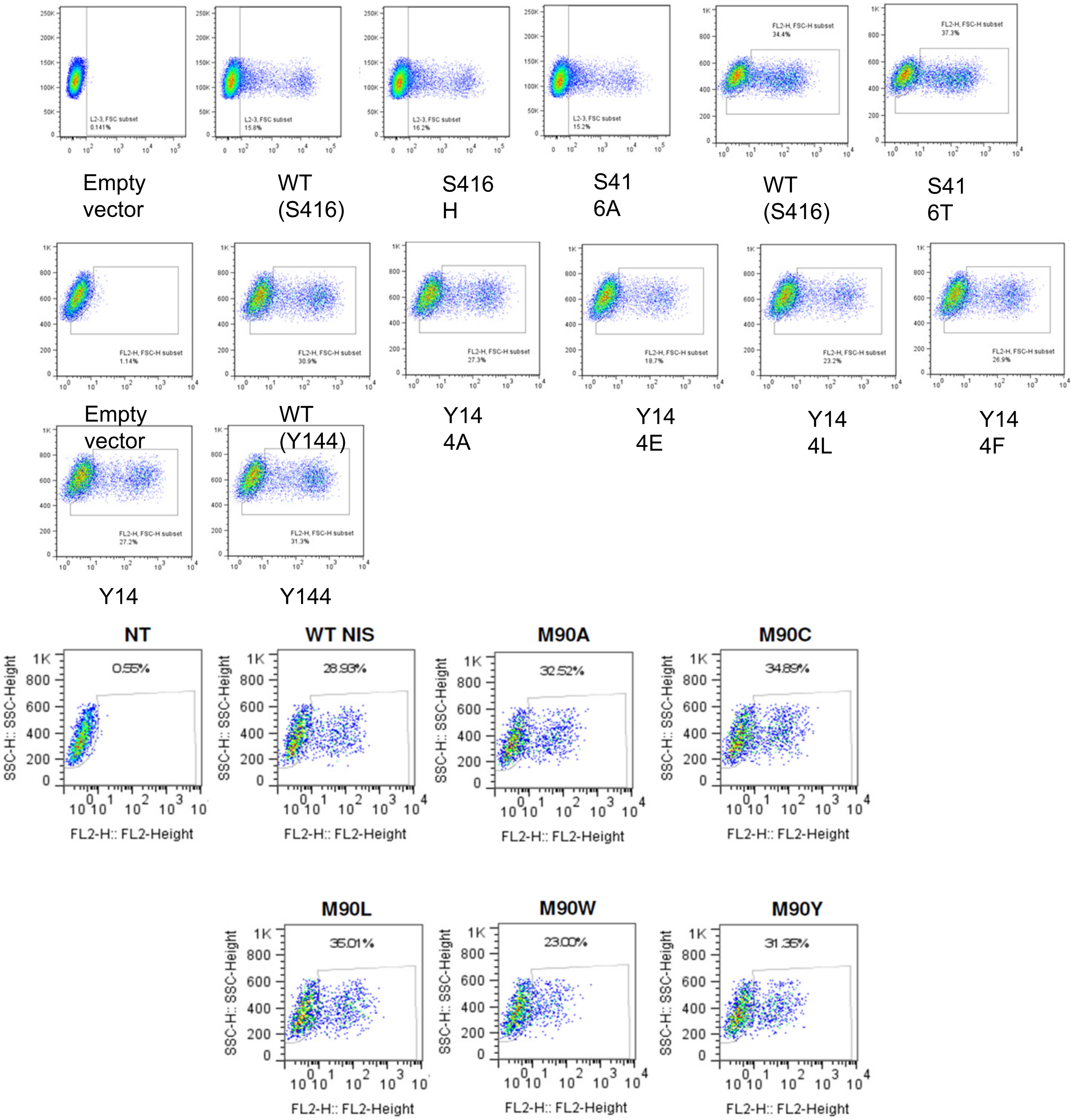

Empty vector
WT (S416)
S416 H
S41 6A
WT (S416)
S41 6T
Empty vector
WT (Y144)
Y14 4A
Y14 4E
Y14 4L
Y14 4F
Y14
Y144
NT
0.55%
WT NIS
28.93%
M90A
32.52%
M90C
34.89%
M90L
35.01%
M90W
23.00%
M90Y
31.35%
SSC-H:: SSC-Height
FL2-H:: FL2-Height

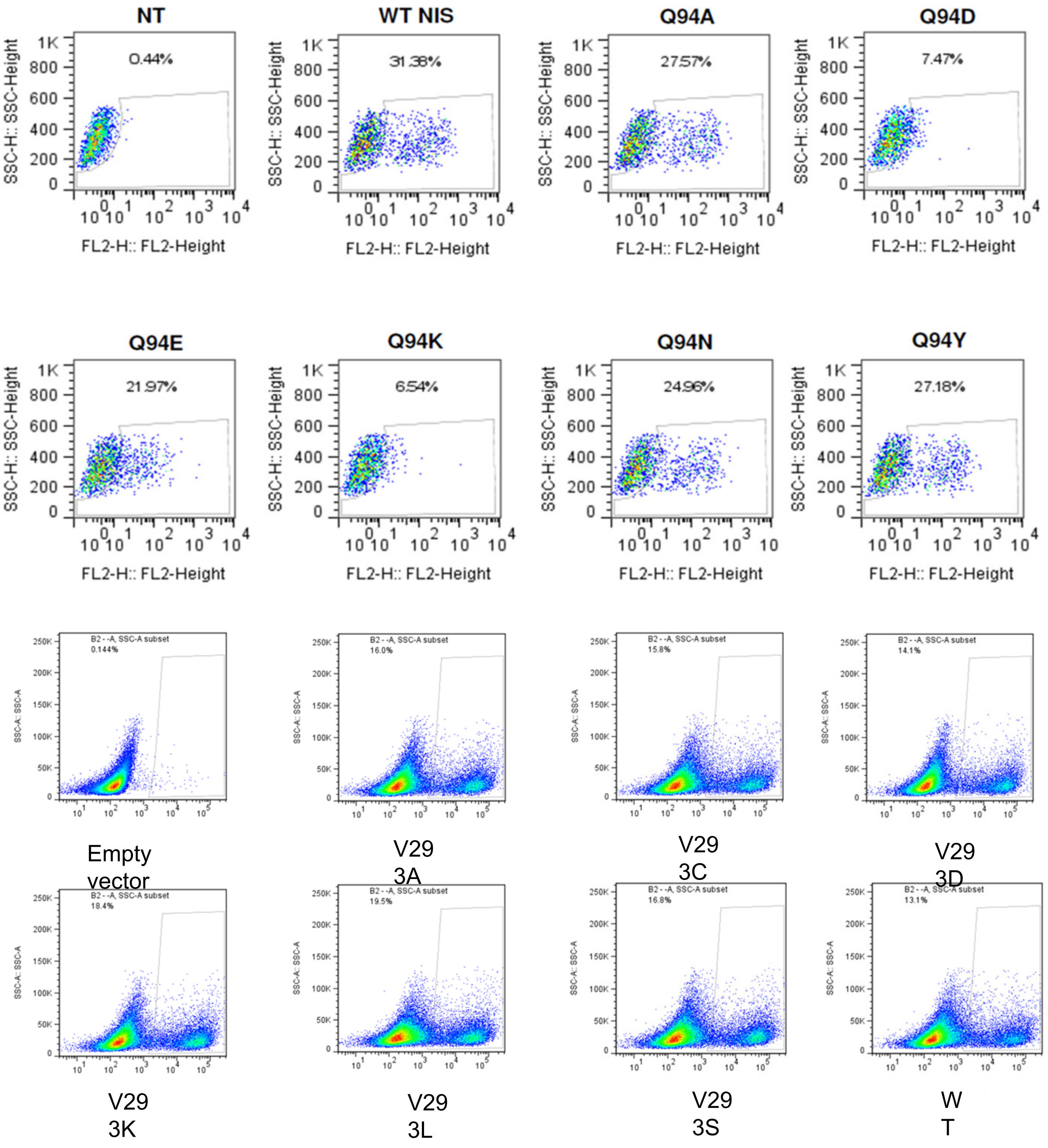


**Supplementary Fig. 4 NIS mutant proteins are properly targeted to the plasma membrane.** Nontransfected HEK cells (NT) and HEK cells transfected with WT NIS or the indicated NIS mutants were incubated under nonpermeabilized conditions with an anti-HA antibody that recognizes the extracellular N-terminus HA epitope and analyzed by flow cytometry. The *x* axes show the intensity of the fluorescence of each single cell; the *y* axes the values of the side scatter parameters. For each experiment, NT cells were used as a reference to identify the negative cells and determine the percentage of cells expressing WT NIS or the respective mutants.

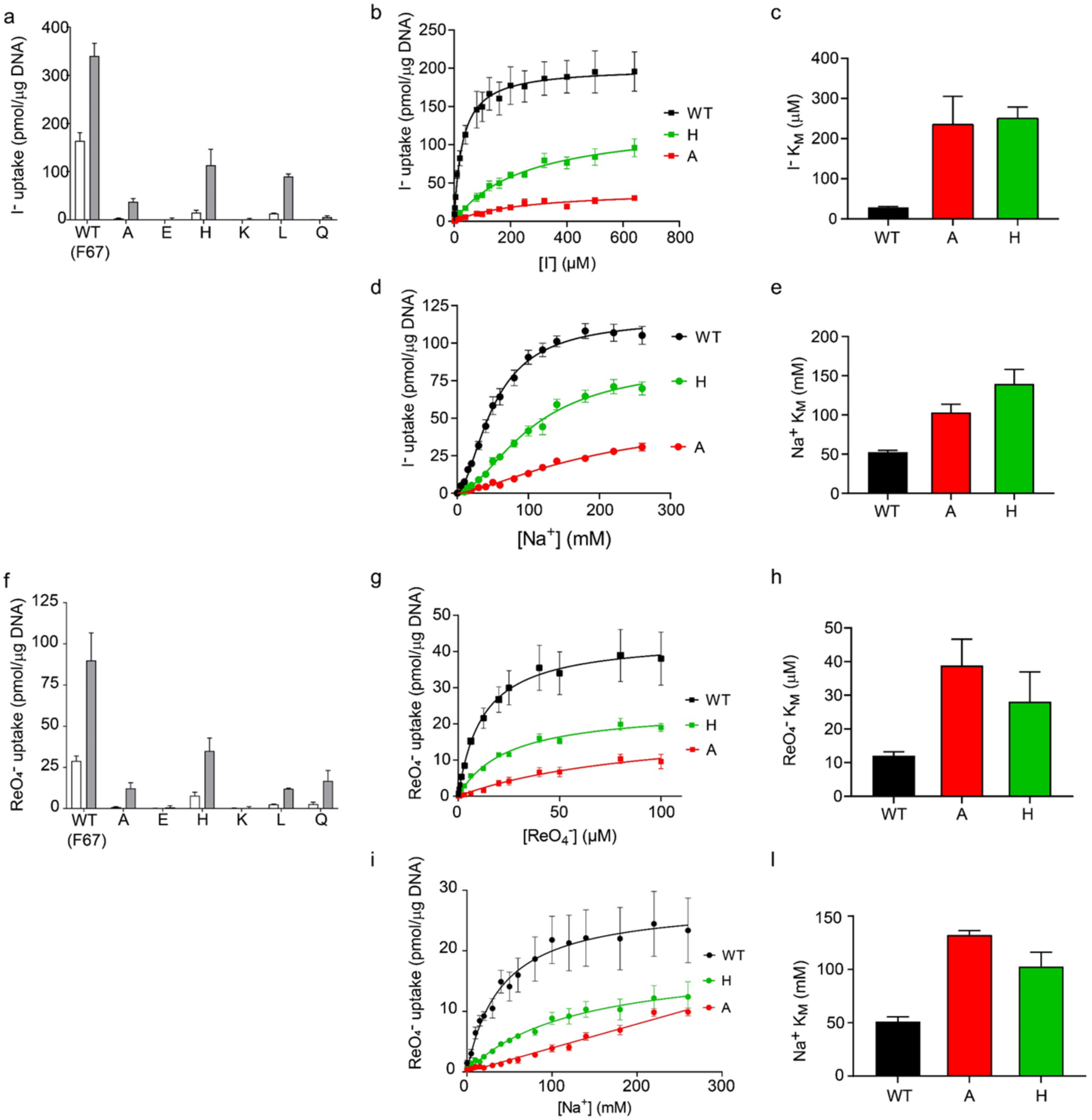


**Supplementary Fig. 5 Effect of single amino acid substitutions at position 67 on $I^-$ and $ReO_4^-$ transport. a.** NIS-mediated $I^-$ uptake at steady state. cDNA constructs coding for NIS mutants in which F67 is replaced with the residues indicated were transfected into COS7 or HEK cells. $I^-$ uptake by these NIS mutants was measured at 20 μM (white bars) and 200 μM (gray bars) $I^-$ at 140 mM $Na^+$ for 30 min with or without the NIS-specific inhibitor $ClO_4^-$ (values obtained in the presence of $ClO_4^-$, which are < 10% of the values obtained in its absence, have already been subtracted). Results are given as pmols of $I^-$ accumulated/μg DNA ± SE. Values represent averages of the results from two or three different experiments, each of which was carried out in triplicate. **b and d**. Kinetic analysis of initial rates of $I^-$ uptake (2-min time points) determined at 140 mM $Na^+$ and varying concentrations of $I^-$ (b), and at varying concentrations of $Na^+$ and 500 μM $I^-$ (d). **c and e.** $I^-$ $K_M$ and $Na^+$ $K_M$ values determined from b and d, respectively. **f.** NIS-mediated $ReO_4^-$ uptake at steady state. $ReO_4^-$ uptake by these NIS mutants was measured at 3 μM (white bars) and 30 μM (gray bars) $ReO_4^-$ at 140 mM $Na^+$ for 30 min with or without the NIS-specific inhibitor $ClO_4^-$ (values obtained in the presence of $ClO_4^-$, which are < 10% of the values obtained in its absence, have already been subtracted). Results are given as pmols of $I^-$ accumulated/μg DNA ± SE. Values represent averages of the results from two or three different experiments, each of which was carried out in triplicate. **g and i**. Kinetic analysis of initial rates of $ReO_4^-$ uptake (2-min time points) determined at 140 mM $Na^+$ and varying concentrations of $ReO_4^-$ (g), and at varying concentrations of $Na^+$ and 100 μM $ReO_4^-$ (i). **h and j.** $ReO_4^-$ and $Na^+$ $K_M$s values determined from g and i, respectively.

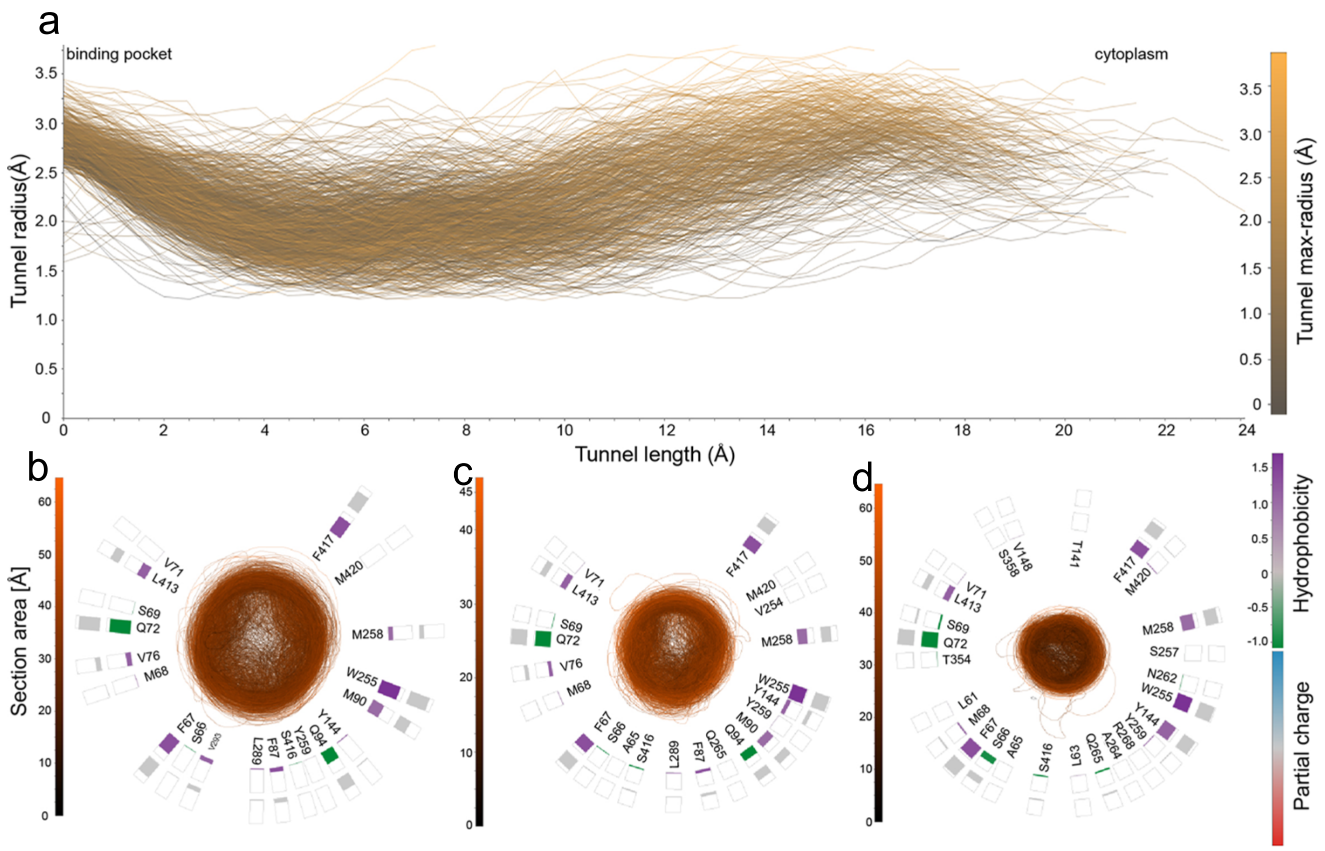


**Supplementary Fig. 6 Pathway analysis from NIS-I⁻ MD simulation snapshots. a.** Cluster of tunnels identified in 1000 snapshots from the MD simulations carried out with NIS-I⁻. **b-d.** section of the tunnel cluster taken respectively at 3.5 Å (a), 5 Å (b), and 10 Å (c) from the position of I⁻ in the structure.

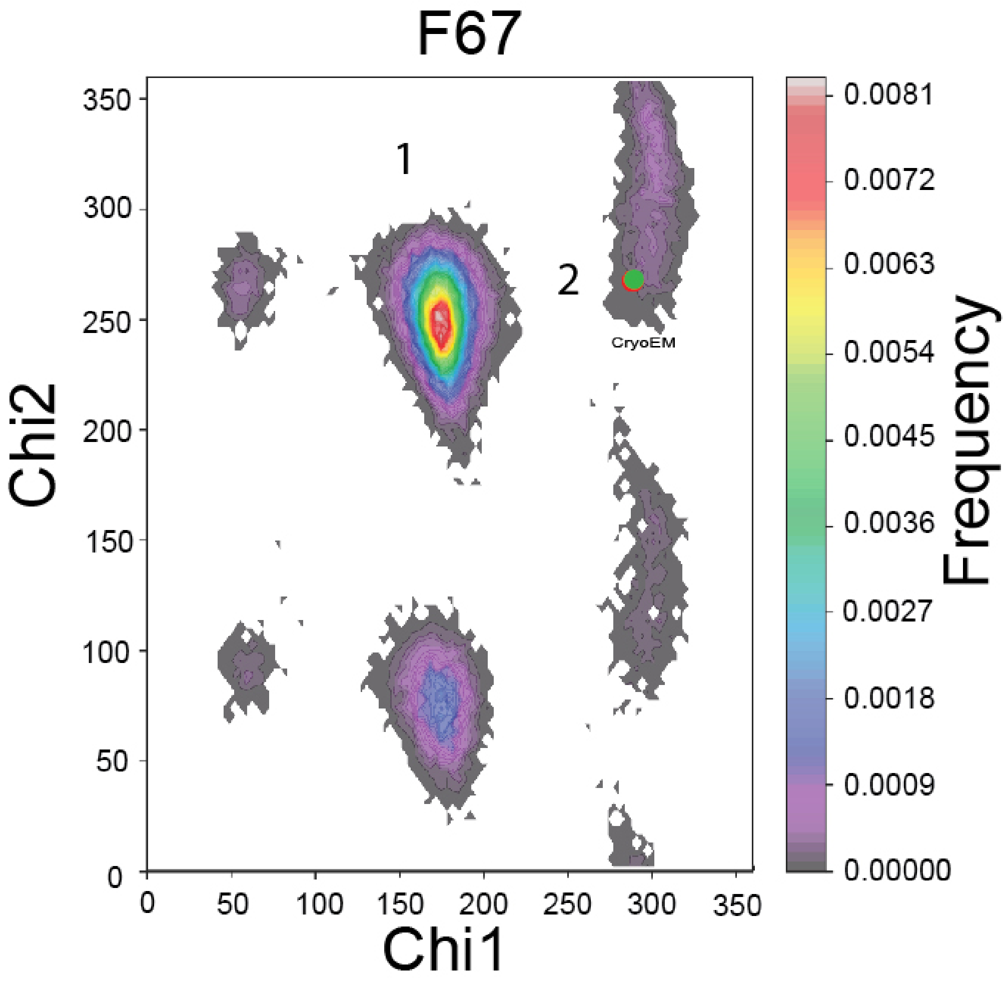


**Supplementary Fig. 7 F67 conformations open up the ion pocket toward the exit pathway.** Ramachandran plot of the chi1 and chi2 side chain dihedral angles of F67 visited during the MD simulations with NIS-I$^-$. The dihedral angles selected are the principal determinants of the position of the side chain. The excursions of these dihedral angles (during the MD simulations) away from the conformational basins corresponding to the cryoEM structure (green dot in basin 2) and toward conformational basins (blue dot in basins 1) open up the exit path. In these histograms, the frequency of a given conformational state is indicated by a rainbow gradient from deep purple (0 frequency) to red (highest frequency).

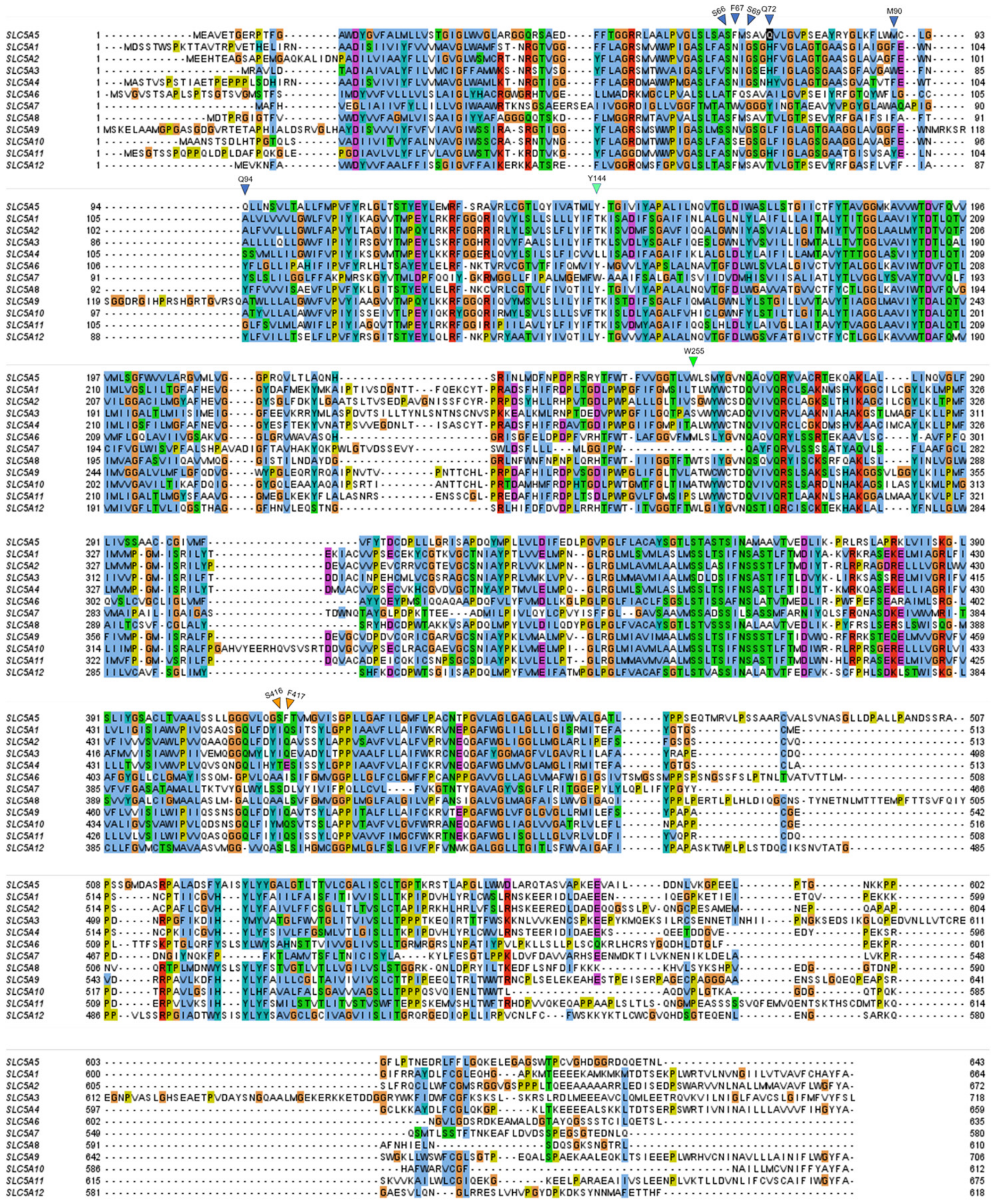


**Supplementary Fig. 8 | Alignment of the sequences of the members of the SLC5 family.**
